# A novel reusable transcriptome-wide association study workflow used to map key genes linked to important cattle traits

**DOI:** 10.1101/2025.06.10.658680

**Authors:** S. Jayaraman, P. Krishna Chitneedi, N. Kumar Kadri, G. Costa-Monteiro-Moreira, M. Salavati, C. Charlier, D. Boichard, M-P. Sanchez, H. Pausch, C. Kühn, J.G.D. Prendergast, E.L. Clark

## Abstract

Transcriptome-wide association studies (TWAS) are a powerful approach for studying the genes underlying complex traits by directly integrating GWAS and gene expression datasets. In cattle, they have been previously applied to identify genes driving fertility, milk production, and health. However, these studies have also highlighted several challenges, from difficulties in reproducing these complex analyses to limitations from poor genotype calls, especially when called directly from RNA sequencing data. To address these and other challenges, for the H2020 BovReg Project, we have developed a streamlined, species-agnostic, and reusable Nextflow TWAS workflow to integrate transcriptomic and GWAS summary statistic datasets. Our workflow first generates accurate genotype calls and gene expression prediction models from transcriptomic datasets and then applies these tools to impute gene expression levels into GWAS cohorts, enabling the association of genes with traits of interest. We explore optimal strategies for calling genetic variants directly from transcriptomic data and illustrate that using imputation approaches specifically designed for low-pass sequencing data can improve variant calling over previously adopted methods. We demonstrate the utility of our TWAS workflow by applying it to both novel and publicly available GWAS cohorts for cattle, detecting novel gene-trait associations for complex traits. Using a new transcriptome annotation of the cattle genome generated for the BovReg project we also illustrate how previously un-assayable associations can be detected. The results and the workflow we present, provide a new resource for the community and contribute to a better understanding of the molecular drivers of complex traits in cattle with the goal of eventually leveraging this information in future breeding decisions.

## Introduction

Many quantitative trait loci (QTL) associated with complex traits in cattle have been discovered using genome-wide association studies (GWAS) [1–3]. The QTL database for cattle (Cattle QTLdb) currently contains 192,247 QTLs/associations (correct as of September 26th, 2024) [4]. However, a major challenge in genetic association studies is determining the biological mechanisms through which a locus is linked to a trait. One approach to achieve this is to characterise which variants in the locus are linked to changes in gene expression. For example, expression QTL (eQTL) analyses can be used to determine which genetic variants in a locus are linked to changes in transcription levels and, more specifically, to identify which genes are involved, providing potential functional mechanisms driving the observed genetic association [5]. Several eQTL studies have been published recently for both *Bos taurus* and *Bos indicus* cattle. For example eQTLs have been generated using blood, milk cells, and mammary gland tissues from Holstein cattle [6–10], whole blood from Brahman cattle [11], muscle from Angus-Brahman crossbreed cattle [12], Limousine [13] and Nellore cattle [14,15], and liver and muscle from Simmental cattle [16]. However, most of these studies only report the broad genomic region of eQTLs and are not directly integrated with GWAS data to identify the putative regulatory variants driving downstream phenotypic differences directly.

Transcriptome-wide association studies (TWAS) are a powerful approach for detecting complex trait-associated genes (reviewed in [17]) by directly integrating GWAS and gene expression datasets [17]. In this approach, eQTL cohorts are used to build predictive models to impute gene expression levels from genotypes. These models can then be applied to independent GWAS cohorts, where genotypes and phenotypes are available, but expression data is not. In this way the imputed gene expression values can be directly tested for an association with the downstream traits, allowing researchers to directly link genes, rather than genetic variants, to phenotypes [18].

Recent studies, including the Cattle Genotype–Tissue Expression atlas (CattleGTEx) project [19], have demonstrated the power of TWAS in identifying genes driving complex traits, including lactation [7], male fertility [20], female fertility [11], health [21,22], and carcass traits [12]. However, these studies have also highlighted a number of challenges. The full TWAS workflow requires a large number of steps and can therefore be challenging to implement from scratch. Additionally, many RNA-seq cohorts lack matching genotype data. To address this, the CattleGTEx project successfully adopted an approach of calling genetic variants from the RNA-seq data directly, but as demonstrated recently by Leonard et al. (2024), the primary limitations of such an approach are the limited coverage outside coding regions and allele specific expression levels confounding genotype calls [23]. Likewise, CattleGTEx and others have combined disparate RNA-seq cohorts together that may introduce issues such as batch effects. Finally, limitations of the cattle genome annotation in comparison to other species such as humans, can mean that gene-trait or transcript-trait associations can be missed.

To address these challenges, we have developed a streamlined, species-agnostic, and reusable Nextflow TWAS framework to integrate transcriptomic data and GWAS summary statistics. Our TWAS workflow integrates available tools including GLIMPSE [24], STAR [25], PrediXcan [26], and MetaXcan [27] to both generate gene expression prediction models from transcriptomic datasets, and to use these models to impute gene expression levels into GWAS cohorts to associate genes with traits of interest. As part of this, we explore optimal strategies for calling genetic variants directly from RNA-seq data and illustrate that using imputation approaches specifically designed for low-pass sequencing data can improve variant calling over previously adopted methods. We demonstrate the utility of this workflow by applying it to both novel and publicly available [28] GWAS cohorts spanning a total of 21 unique traits, and illustrate how it is able to detect novel gene-trait associations. Using a new transcriptome annotation of the cattle genome, generated by BovReg Project partners, from three different populations of cattle across a wide diversity of tissues (described in [29]) we also illustrate how previously un-assayable associations can be detected.

The results and workflow we provide are important resources to help improve our understanding of the molecular drivers of complex traits in cattle populations, with the goal of eventually leveraging this information in breeding decisions.

## Results

### A Nextflow workflow for performing TWAS

To make TWAS tractable for the community, we have developed a Nextflow workflow that spans the whole set of analyses: from the processing of raw fastq data (including datasets downloaded from the public repositories where relevant), through read mapping, variant calling, expression quantification, TWAS model generation and finally TWAS. This workflow is summarized in Figures 1A and B.

**Figure 1:**
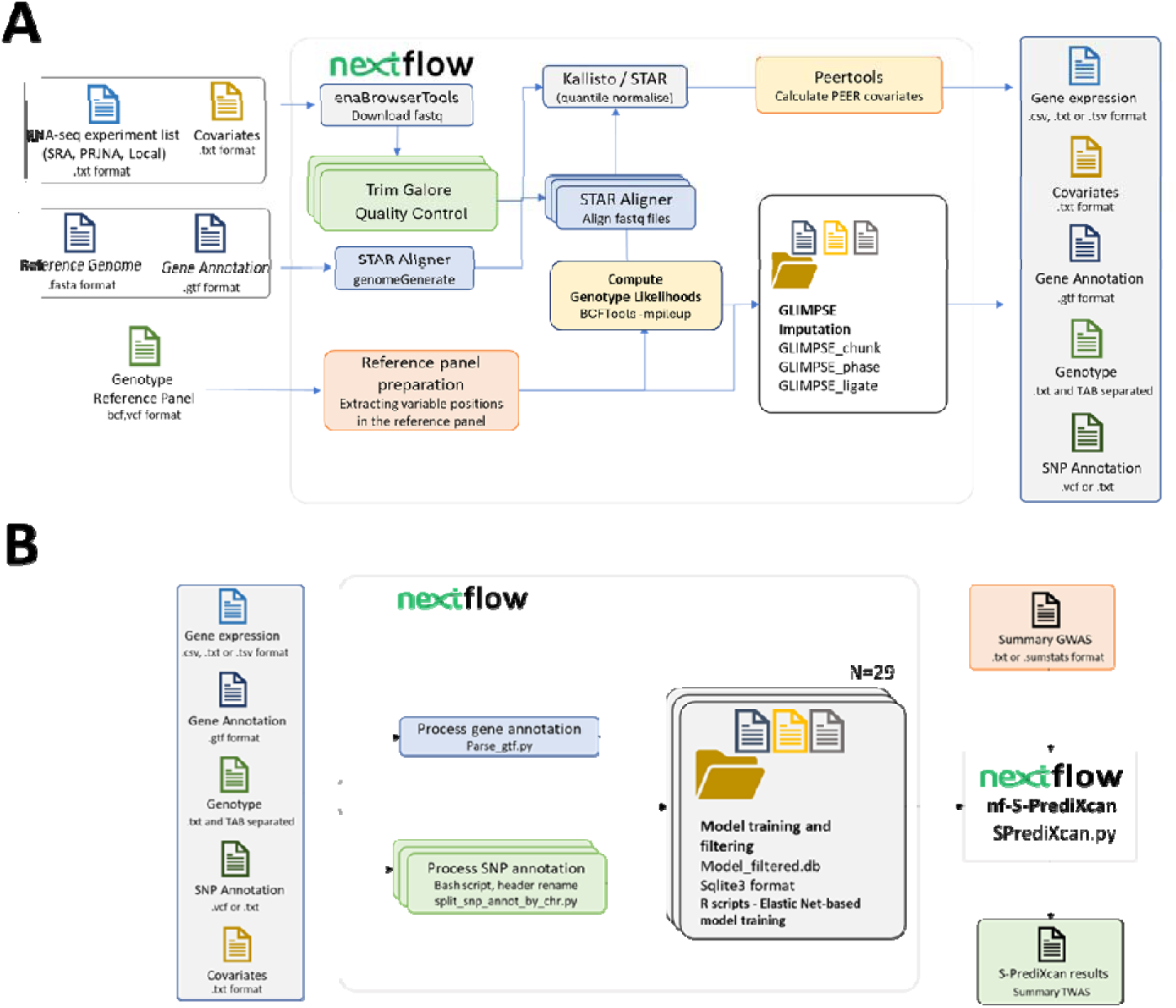
Schematic of the pipeline to generate a PredictDB database compatible with MetaXcan from samples without paired genotype datasets using just RNA-seq datasets. A) Detailed workflow of the initial steps in the TWAS pipeline. B) S-PrediXcan TWAS workflow using the outputs from the pipeline shown in A.

Novelties of this workflow include that it allows researchers with their own suitable datasets to train and run TWAS using our scripts totally in-house. Importantly the pipeline is flexible. For example, in instances where a user does not have their own cohort of matching RNA-seq and genotypes, the workflow provides an option to call genotypes directly from their own or publicly available RNA-seq datasets. Further advantages include that this workflow also allows consortia to undertake consistent TWAS approach without the need to share sensitive individual genotype and phenotype information that can be a constraint on such analyses. Consequently, we have developed a streamlined, reusable pipeline for TWAS in any species, that is applicable even where a user lacks their own suitable RNA-seq datasets.

### GLIMPSE provides improved variant calls from RNA-seq data

As part of the development of this workflow, we assessed alternative analysis approaches at each step. One of the most critical steps is generating a dataset of gene expression and matching genotype calls. However, in livestock few large cohorts of RNA-seq data with matching genotype array or genome sequencing data exist. Consequently, calling and imputing variants directly from RNA-seq data has been adopted to overcome this limitation. For example, in the CattleGTEx project [19], variants were called and imputed from RNA-seq datasets using GATK [30] and BEAGLE [31] respectively. However, across most of the genome RNA-seq reads are perhaps more akin to low-pass sequencing data, in that few if any reads are found covering most bases. We therefore tested the relative performance of using GLIMPSE, a tool specifically designed for low coverage imputation [32], to call and impute variants from three Holstein-Friesian (*Bos taurus taurus*) and three Nellore (*Bos taurus indicus*) samples for which we had both immune cell RNA-seq data and high quality variant calls from matching ∼30X coverage whole genome sequencing (WGS) data [33]. As shown in Figure 2 imputation with both GLIMPSE and BEAGLE showed substantial improvements in genotype concordance with the WGS data over the raw RNA-seq data variant calls. However, GLIMPSE generally not only calls more genotypes but does so more accurately than BEAGLE. This is observed across read depths (Figure 2) and genotype quality cutoffs (Figure 3) suggesting GLIMPSE is the preferred approach for such studies and is consequently the chosen option in our workflow.

**Figure 2:**
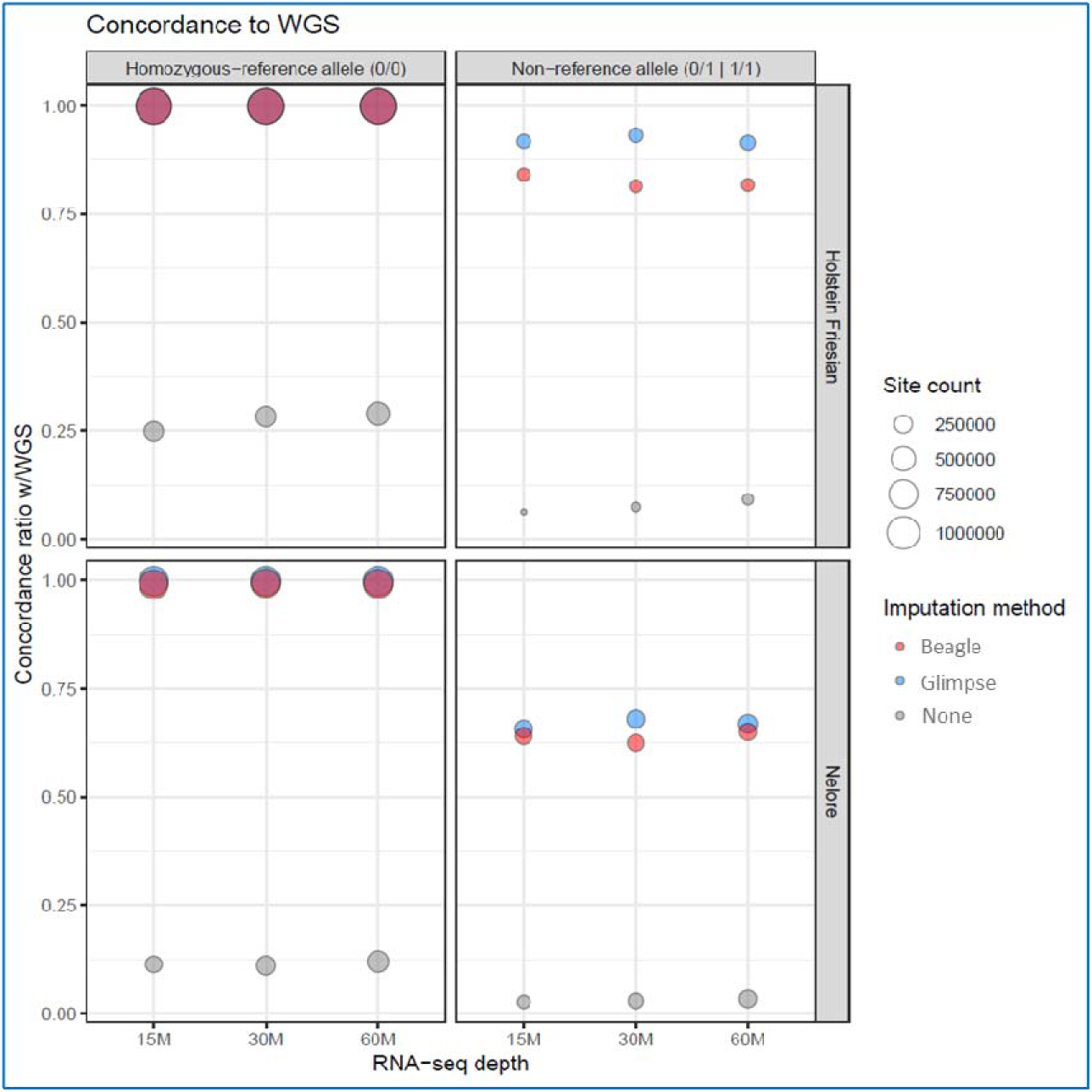
Concordance to whole-genome sequencing (WGS) for variant calling using different imputation methods across RNA-seq depths. The proportion of genotypes that match the high coverage WGS calls is shown for homozygous-reference (0/0, left panels) and non-homozygous reference calls (0/1 and 1/1, right panels) in Holstein Friesian (top row) and Nellore cattle (bottom row). RNA-seq read counts of 15M, 30M, and 60M reads are compared on the x-axis. Imputation methods include BEAGLE (red), GLIMPSE (blue), and non-imputed mpileup calls (grey). The size of each point corresponds to the number of sites evaluated.

**Figure 3:**
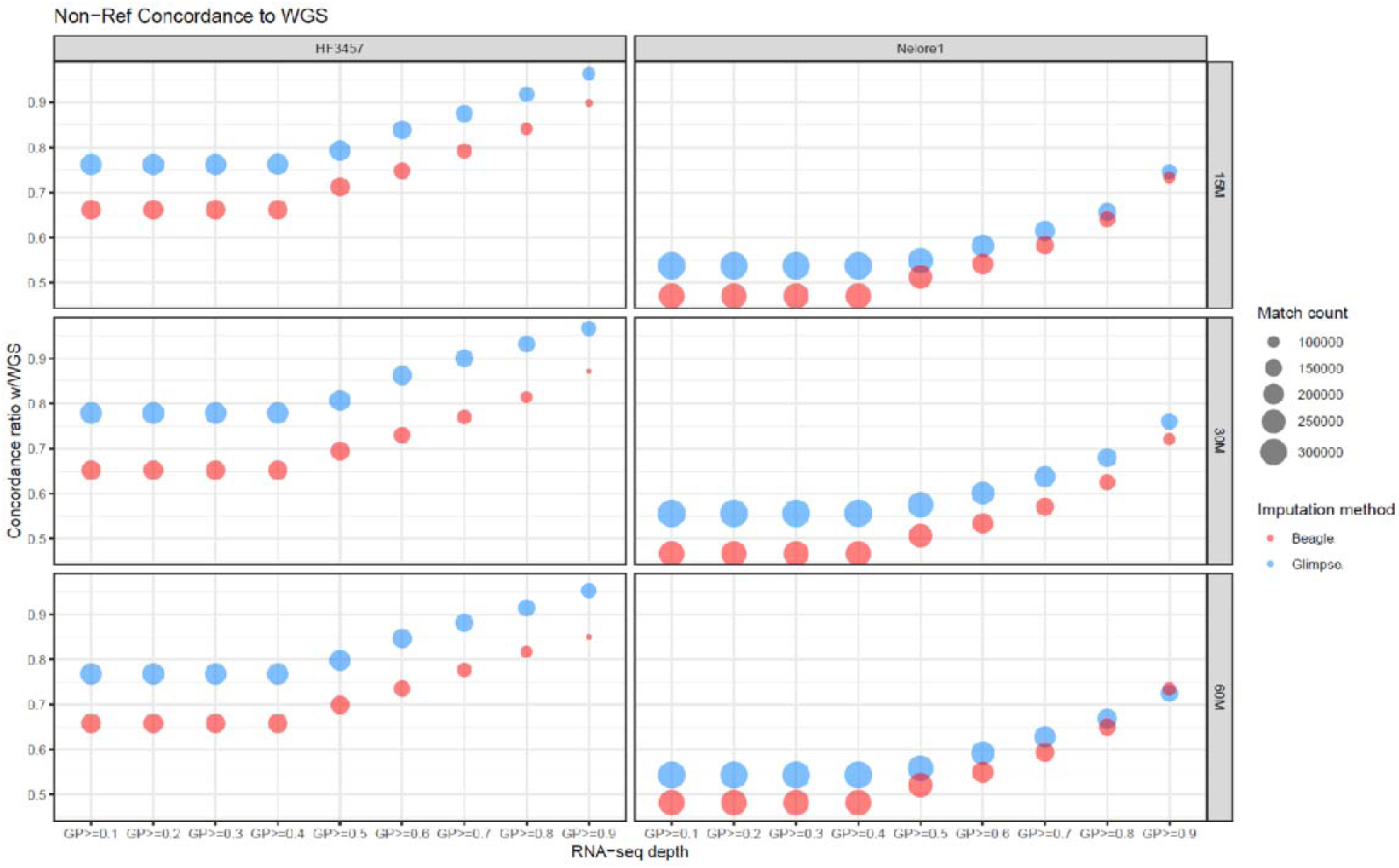
Non-reference Concordance to whole-genome Sequencing (WGS) across RNA-seq depths and genotype probabilities (GP). The proportion of non-reference alleles (0/1 and 1/1) matching WGS is shown for Holstein Friesian (*HF3457*, left panels) and Nellore (*Nellore1*, right panels) cattle at RNA-seq depths of 15M, 30M, and 60M reads (rows). The x-axis represents increasing genotype probability (GP) thresholds. Imputation methods include BEAGLE (red) and GLIMPSE (blue). The size of each point corresponds to the number of matched genotypes. The proportion of genotypes matching WGS improves with increasing GP thresholds and RNA-seq depth, with notable differences in performance between the imputation methods.

It should be noted that both approaches performed comparatively poorly on the *Bos indicus* samples, for rarer variants (Supplementary Figure 1). This likely reflects the bias of the 1000 Bull Genomes imputation cohort [34] towards European *Bos taurus* breeds and highlights the importance of using a more representative reference cohort if imputing sequence variant genotypes in this sub-species.

### RNA-Seq data used for TWAS model training

To further test the workflow, we downloaded RNA-seq data from public repositories for five distinct, cattle tissues: muscle, rumen, mammary gland, liver and jejunum. To minimize batch effects, and ensure sufficient power, we restricted these data to RNA-seq cohorts with a minimum of 48 samples (Table 1). These samples were then processed through our workflow, involving read processing and QC using Trim Galore, genome alignment and gene quantification using STAR [25], transcript quantification using Kallisto [35], covariate estimation using PEER tools [36], variant calling using bcftools [37] and genotype imputation using GLIMPSE [32] (Figure 1A). See methods section for more details.

**Table 1:** Details of RNA-seq datasets downloaded from the public repositories that were used as RNA-Seq cohorts for TWAS model training.

| Bioproject | Breed | Tissue | No. of Samples | Reference |
| --- | --- | --- | --- | --- |
| PRJNA682457 | Holstein | Lactating Mammary Gland | 145 | [38] |
| PRJEB34570 | Charolais x Holstein | Jejunal mucosa | 66 | [39] |
| PRJEB34570 | Charolais x Holstein | Liver | 48 | [39] |
| PRJEB34570 | Charolais x Holstein | Skeletal muscle | 48 | [39] |
| PRJEB34570 | Charolais x Holstein | Rumen | 63 | [39] |

To check for any batch effects or anomalous samples in these datasets we conducted a principal component analysis (PCA) to cluster samples by their gene expression levels. As shown in Figure 4 the samples clustered closely together by tissue type, with the two gastro-intestinal tissues, jejunum and rumen, showing the closest association of any of the tissues, reflecting similarities in their physiology and function.

**Figure 4:**
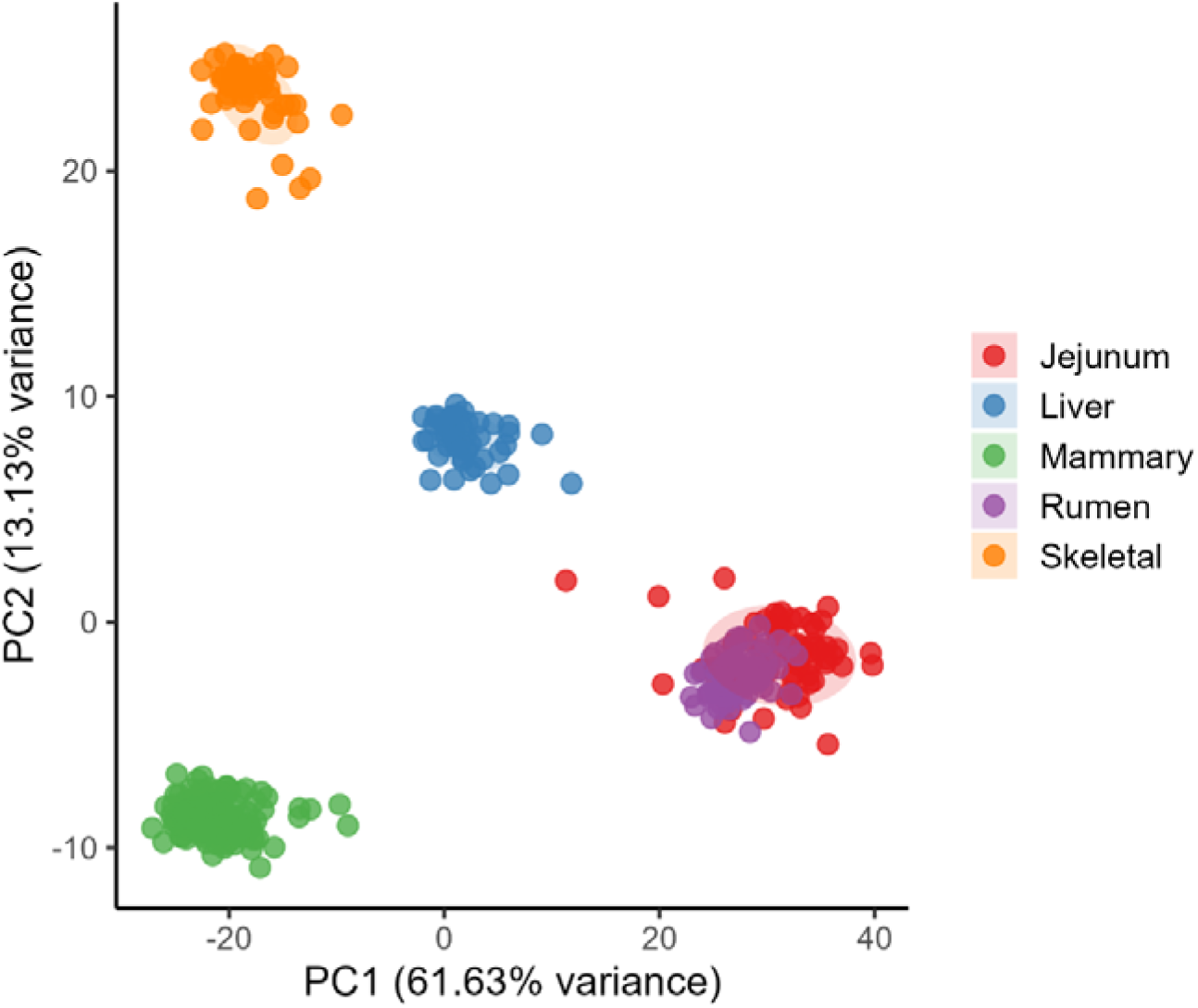
Principal Component Analysis (PCA) of Gene Expression Across Tissues. Principal components (PC1 and PC2) are plotted to visualize the variance in gene expression profiles across five tissues (Table 1): Jejunum (red), Liver (blue), mammary gland (green), rumen (purple), and skeletal muscle (orange). Distinct clustering of samples by tissue type indicates clear tissue-specific expression patterns.

### Increased mapping rate to annotated features in new bovine genome annotation

The BovReg consortium has recently generated an updated gene and transcript annotation of the cattle genome, the version used for the analysis presented in this manuscript is BovReg_RNA-Seq_CAGE_merged_track_sorted_LIEGE.gtf (Costa-Monteiro-Moreira et al. https://figshare.com/articles/dataset/BovReg_TWAS_Annotation_File_BovReg_RNA-Seq_CAGE_merged_track_sorted_LIEGE_gtf/29264816?file=55229522). The pipeline used to generate the new annotation is available via the BovReg GitHub repository (https://github.com/BovReg/rnaseq). This new annotation was generated from PolyA+ RNA, total RNA, small RNA and CAGE data from a diverse set of 24 tissue types from six individual cattle of different breeds, and sexes. The samples are described in Salavati et al. [29], the six individuals belonged to three populations representing dairy (Holstein Friesian; HF), beef–dairy cross (Charolais × Holstein F2; CH), and beef composite (Kinsella composite; KC) cattle. We explored the relative merits of using this annotation set relative to that available via the Ensembl genome browser (ARS-UCD1.2 Ensembl v106) and found that 14,975 gene models (and 30,830 transcript models) were novel and unique to BovReg. As shown in Figure 5, use of the BovReg annotation substantially improved RNA-seq mapping rates over the ARS-UCD1.2 Ensembl v108 cattle genome annotation. For example, on average almost 70% of RNA-seq reads from our cohorts mapped to a BovReg gene annotation compared to only 60% aligning to an Ensembl gene. BovReg annotations were consequently used in downstream analyses.

**Figure 5:**
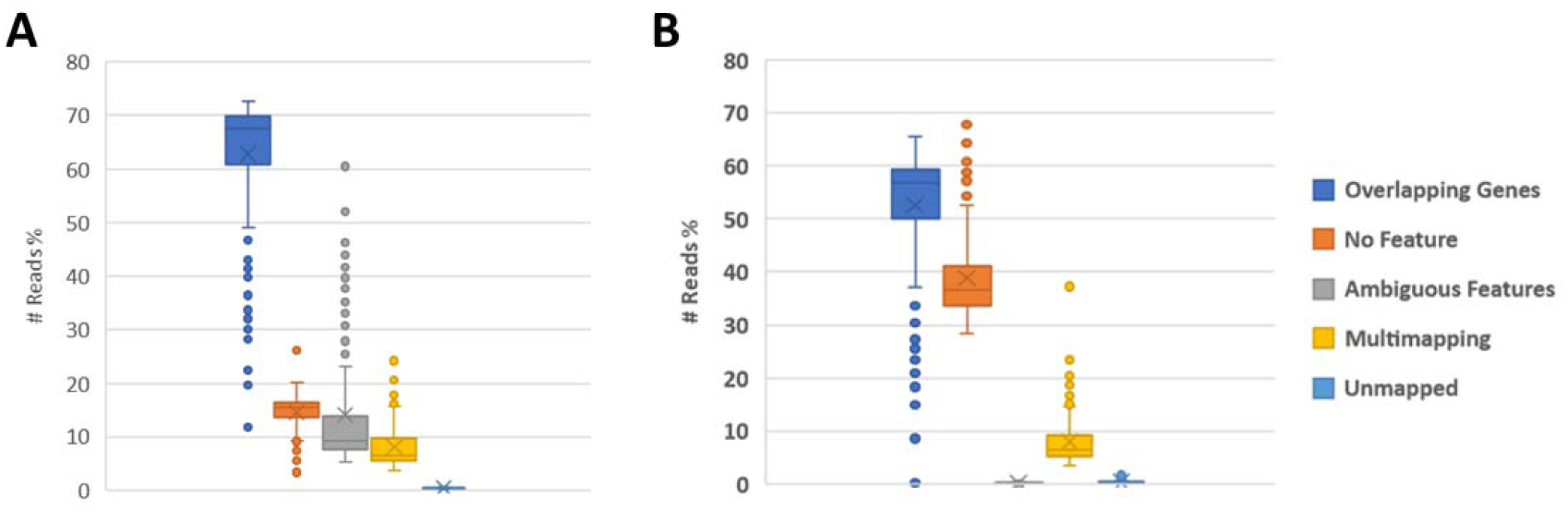
Comparison between feature count after aligning RNA-seq reads with STAR using A) BovReg vs B) Ensembl.

### Performance of the workflow for TWAS model training

To demonstrate the utility of any developed TWAS models at providing novel insights into the genes potentially underlying cattle phenotypes we collated summary statistics from four GWAS cohorts, three from BovReg partners, and one publicly available dataset [28] that had tested genetic associations with the phenotypes summarized in Supplementary Table 1. The sizes of these GWAS cohorts ranged from 3000-27000 animals (Supplementary Table 1) and the number of variants tested in each cohort ranged from 1,412,041 to 27,214,879.

Using the GLIMPSE derived genotypes and our novel workflow (Figure 1A) we trained PredictDB TWAS models for predicting gene and transcript levels from the cis-genotypes available in each of the GWAS cohorts (Table 2).

**Table 2:** Table summarizing the total number of variants used in model training for each GWAS cohort paired with tissue-level expression data. The columns represent the GWAS cohorts from various institutions; ETH: Eidgenössische Technische Hochschule Zürich (Swiss Federal Institute of Technology in Zurich), FBN: Forschungsinstitut für die Biologie landwirtschaftlicher Nutztiere (Leibniz Institute for Farm Animal Biology), INRAE: Institut National de Recherche pour l’Agriculture, l’Alimentation et l’Environnement (French National Research Institute for Agriculture, Food and Environment), USDA: United States Department of Agriculture

| Tissue / Variant count for model training | ETH | FBN | INRAE | USDA |
| --- | --- | --- | --- | --- |
| PRJEB34570_jejunum | 8,446,450 | 8,847,999 | 10,657,308 | 1,055,378 |
| PRJEB34570_liver | 8,213,410 | 8,613,633 | 10,423,675 | 1,023,103 |
| PRJEB34570_rumen | 8,394,303 | 8,792,940 | 10,602,733 | 1,052,929 |
| PRJEB34570_skeletal-muscle | 8,540,114 | 8,940,090 | 10,749,162 | 1,069,161 |
| PRJNA682457_lactating_mammary_gland | 9,607,151 | 10,012,564 | 11,812,700 | 1,211,657 |

Figure 6 illustrates the performance of models trained using variants from the USDA GWAS cohort to predict gene expression levels. On average, 556 genes and 2,604 transcripts had valid TWAS models (rho-average > 0.1, p-value < 0.05; see Supplementary Table 2) that met the criteria for downstream analyses. The proportion of suitable models refers to the percentage of genes that passed both upstream filtering criteria (e.g., expression level thresholds, availability of genetic variation) and achieved sufficient predictive accuracy in the model training process.

**Figure 6:**
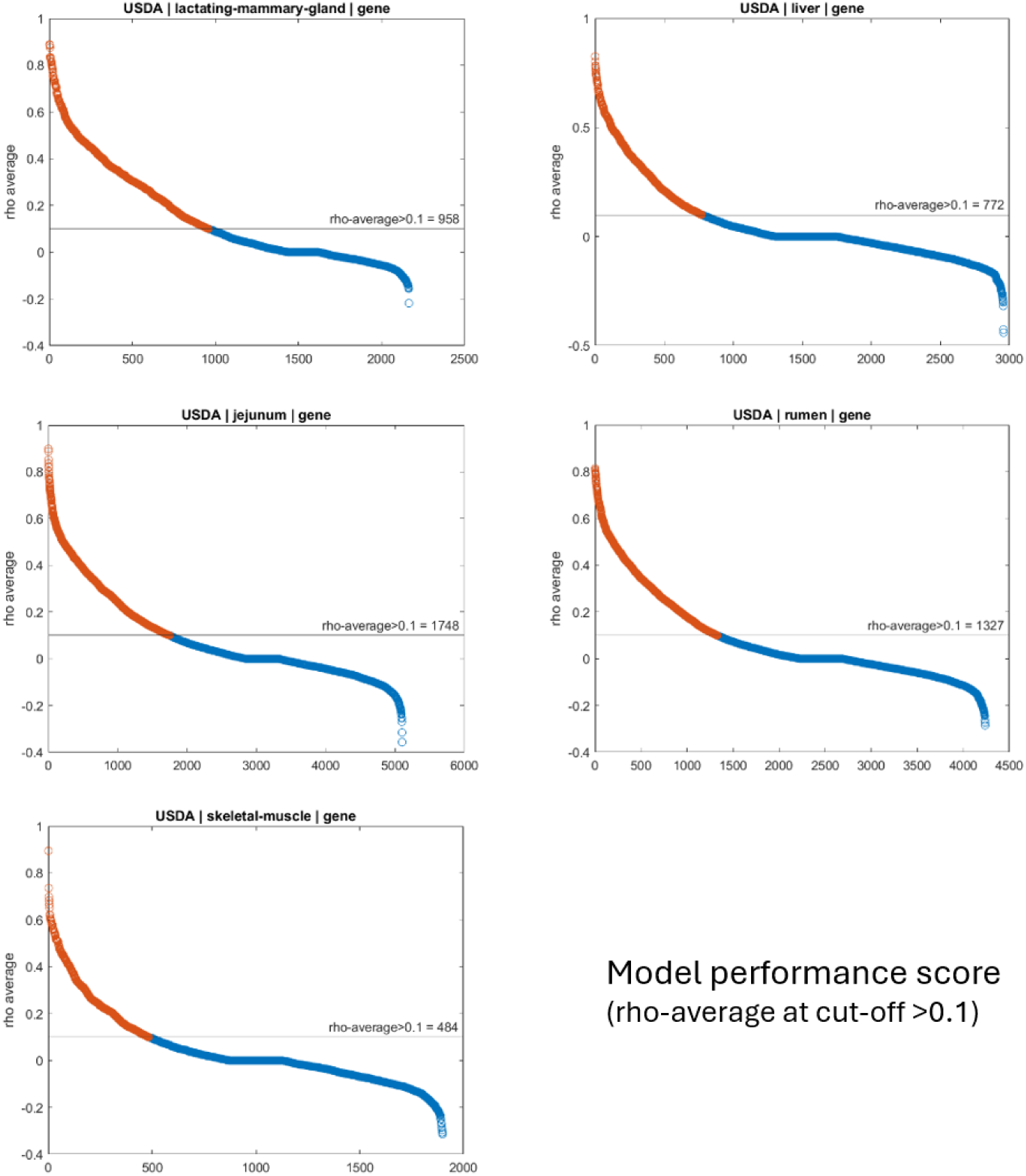
The ability to train predictive TWAS gene-level models using the USDA GWAS cohort variant set. Model performance scores (rho-average) for genes across five tissues: lactating mammary gland, liver, jejunum, rumen, and skeletal muscle. Genes with rho-average > 0.1 (in orange) meet the PredictDB selection criteria, with the number of models exceeding this threshold indicated for each tissue. Blue points represent genes below the threshold.

As expected, the lactating mammary gland tissue had the highest proportion of suitable models, likely due to its larger RNA-seq sample size, which improves the statistical power and robustness of model training. However, despite this, jejunum had the largest absolute number of valid models (937 genes), reflecting the influence of gene expression variability across tissues and the specific filtering applied to the USDA cohort. The relatively low total number of models compared to the number of annotated genes is due to stringent criteria applied during both preprocessing (e.g., genes with low or no expression were excluded) and model evaluation (e.g., requiring rho-average > 0.1 for inclusion).

These results highlight the importance of both RNA-seq sample size and depth for training effective TWAS models. For the USDA GWAS cohort, the relatively limited number of variants (∼1M) used in the analysis (as noted in Supplementary Table 2) may also have contributed to the reduced proportion of suitable models that could be trained.

Figure 7 and 8 and Supplementary figures 2-7 illustrate that few genes or transcripts have a valid model across all of the tissues analysed. For example, only 6 genes and 87 transcripts have a valid model across all five tissues when trained on the set of variants in the USDA GWAS cohort. This partly reflects the fact that some genes and transcripts show tissue restricted expression or less heritability in certain tissues but is also in part driven by the fact that for some tissues fewer valid models were trained due to their sample size and sequencing coverage. This is consistent with results from analyses of human GTEx data that demonstrated that even in their largest cohorts of several hundred samples, not all regulatory variants and genes are detected [40]. In total though 139 to 937 genes and 1031 to 3861 transcripts had a valid model in at least one tissue.

**Figure 7:**
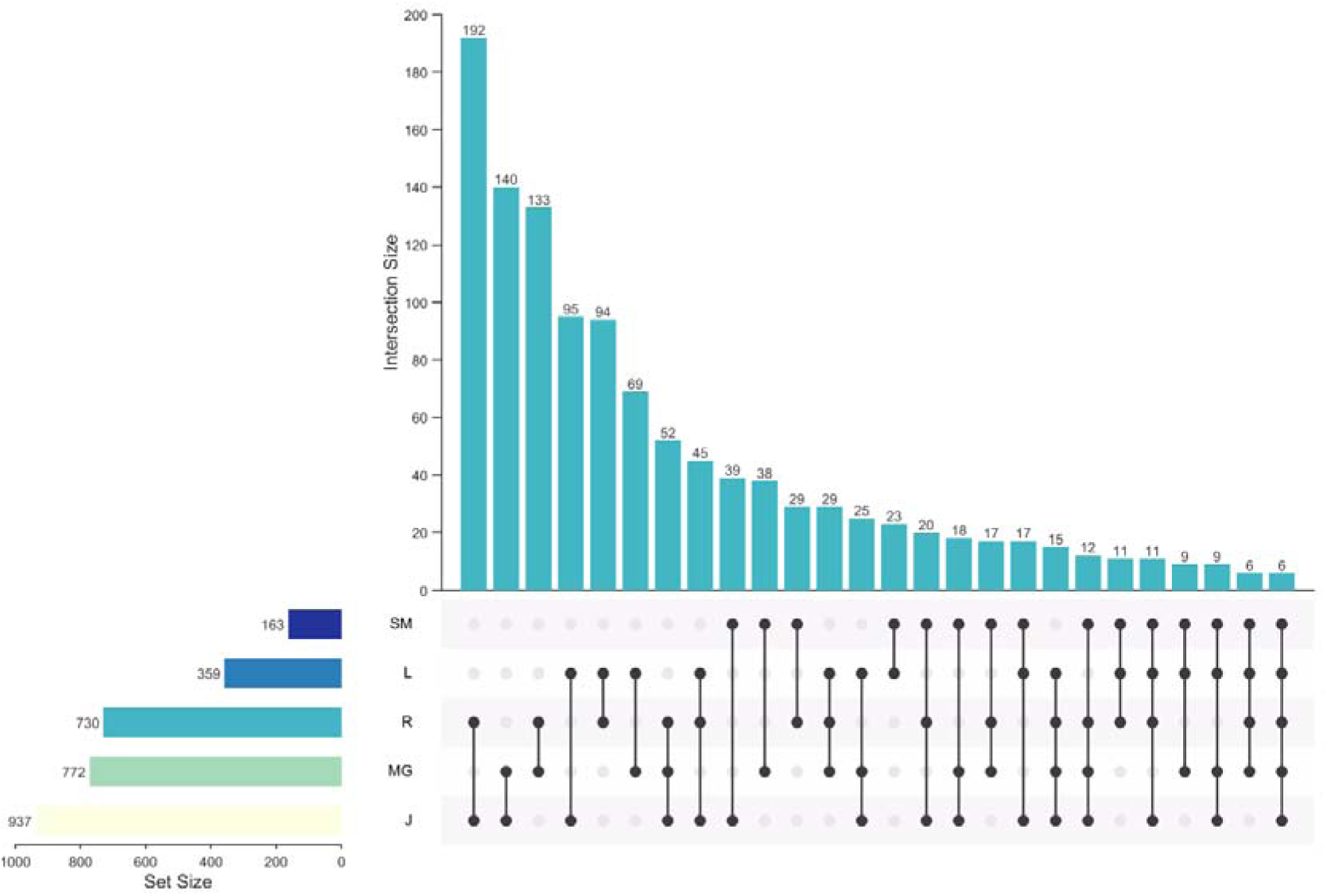
USDA Gene model counts valid across tissues for PredictDB trained models (z score p value <0.05 and rho average >0.1). Tissue names are abbreviated: SM = Skeletal Muscle, L = Liver, R = Rumen, MG = Mammary Gland, J = Jejunum.

**Figure 8:**
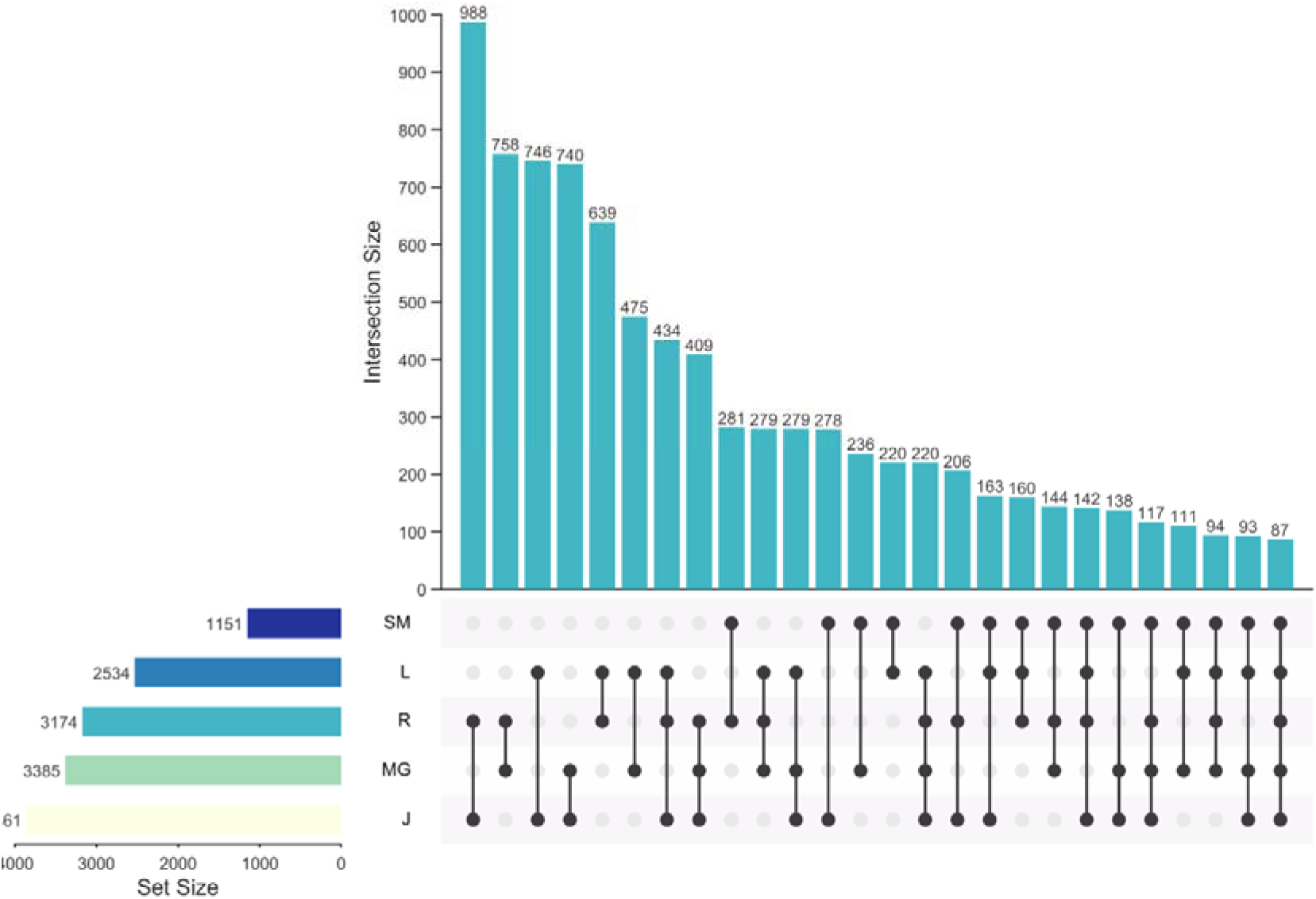
USDA Transcript model counts shared across tissues for PredictDB trained models (z score pvalue <0.05 and rho avg.>0.1). Tissue names are abbreviated: SM = Skeletal Muscle, L = Liver, R = Rumen, MG = Mammary Gland, J = Jejunum.

### Identification of novel trait-gene associations linked to important cattle phenotypes

Using our TWAS models with S-PrediXcan [41] we tested for gene-drivers of the genetic associations seen in the four GWAS cohorts (Supplementary Table 1). To aid in interpretation the novel BovReg gene and transcript models were mapped to Ensembl gene names and annotations where possible (Supplementary Tables 3 and 4). In total, the inferred expression levels of 75 genes and 252 transcripts were significantly associated with at least one trait in at least one tissue. Among these, 6 genes and 23 transcripts were associated with the same trait across tissues, indicating potential involvement in multiple biological contexts. Additionally, 27 genes and 67 transcripts were associated with more than one phenotype, suggestive of potential pleiotropic effects. These values reflect the unique counts of genes and transcripts across studies, accounting for overlap between categories. A detailed study-level breakdown, of unique counts of genes and transcripts across cohorts, accounting for overlap between categories, with the number and list of IDs split by partner/breed/gene-transcript is included in Supplementary Table 5. These results are visualised in Figures 9 and Supplementary figure 8 at the gene level and in Supplementary figures 9 and 10 at the transcript level.

**Figure 9:**
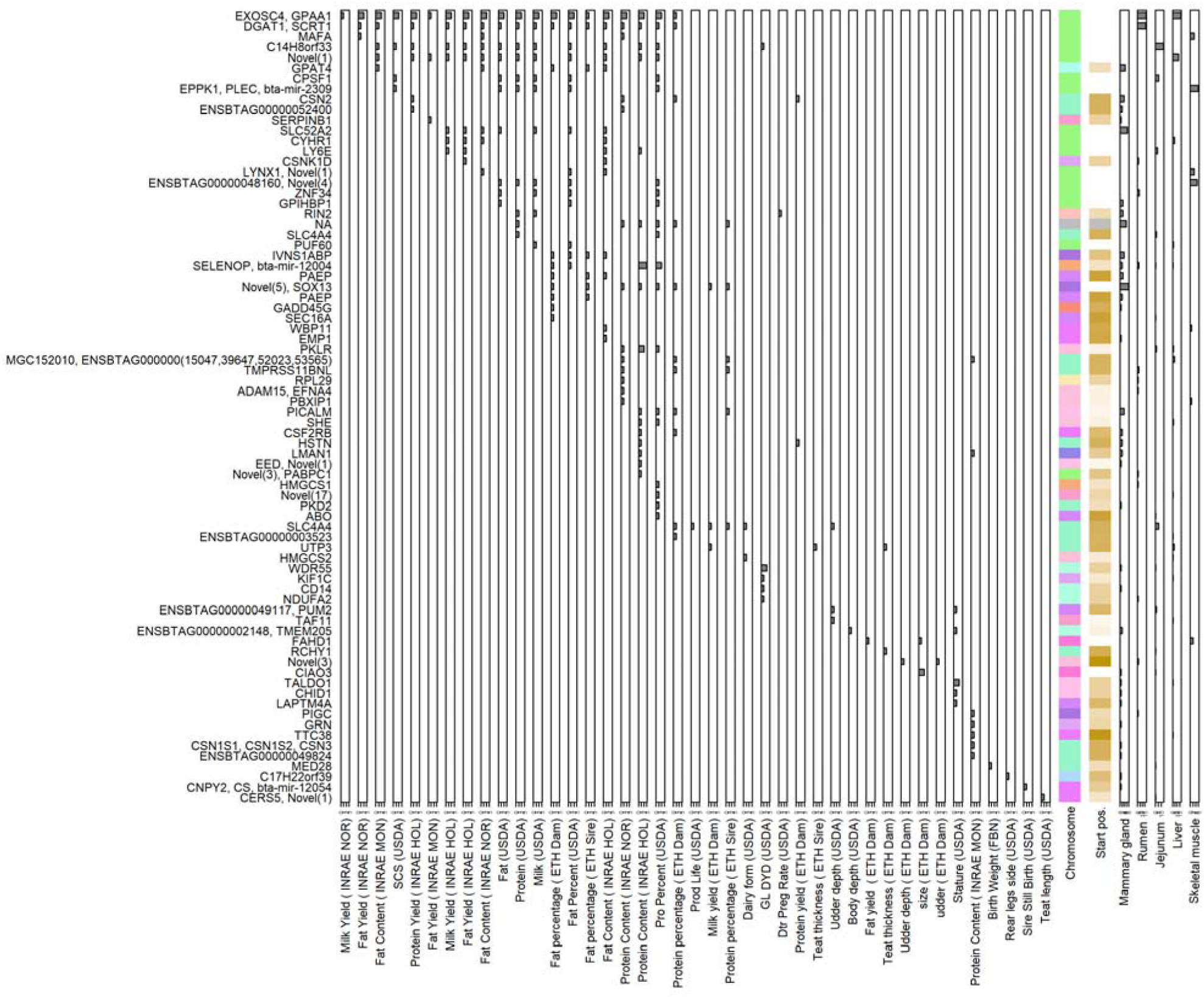
Gene-Trait-Tissue Associations (Bonferroni-adjusted p < 0.05). Bar heights on the furthest left panel indicate the number of tissues in which each gene (Y-axis) is associated with a given trait (X-axis). For each gene, the chromosome and start position are displayed to provide an indication of collocating genes. On the right, results are summarised by tissue with bar heights representing the number of traits the corresponding gene was associated with in the given tissue. This visualization summarizes cross-tissue and cross-trait associations for significant genes.

The top TWAS results for milk yield, protein percentage and stature are shown in Table 3. The full results for all tissues and traits analysed are included in Supplementary Tables 6-8. Our analysis captured many well-known genes linked to, for example, milk production traits in the literature e.g. ABO [28], DGAT1 [42], SLC4A4 [43], PKD2 [44], CSF2RB [45,46] (Table 3).

**Table 3:** The top genes associated with milk yield, protein percentage and stature traits (Bonferroni-adjusted p-value < 0.01), across all four GWAS cohorts. Where a gene/locus is associated with more than one trait and/or in more than one tissue or cohort only the most significant association is shown. “Novel” indicates BovReg transcript models that did not match an Ensembl transcript using gffcompare, with the number in brackets indicating the number of unmatched transcripts. The count within brackets shows the number novel. Note some of the BovReg transcript models span more than one Ensembl transcript. In these cases each gene name is indicated.

| Gene/locus name | Bonferroni-adjusted p-value | Trait | Study | Cohort | Tissue |
| --- | --- | --- | --- | --- | --- |
| DGAT1, SCRT1 | 4.55E-80 | Milk yield | INRAE | HOL | Rumen |
| EXOSC4, GPAA1 | 9.83E-80 | Milk yield | INRAE | HOL | Rumen |
| CPSF1 | 1.97E-75 | Protein percentage | USDA | HOL | Jejunum |
| C14H8orf33 | 7.10E-64 | Milk yield | USDA | HOL | Jejunum |
| Novel(1) | 6.86E-36 | Milk yield | USDA | HOL | Liver |
| ENSBTAG00000048160, Novel(4) | 1.12E-30 | Protein percentage | USDA | HOL | Muscle |
| EPPK1, PLEC, bta-mir-2309 | 2.70E-26 | Milk yield | USDA | HOL | Muscle |
| Novel(5), SOX13 | 4.79E-17 | Protein percentage | ETH | DAM | Mammary Gland |
| SLC4A4 | 8.26E-17 | Protein percentage | USDA | HOL | Jejunum |
| ZNF34 | 1.38E-13 | Protein percentage | USDA | HOL | Rumen |
| HSTN | 1.95E-13 | Protein percentage | INRAE | HOL | Mammary Gland |
| PICALM | 9.36E-13 | Protein percentage | INRAE | HOL | Mammary Gland |
| ENSBTAG00000015047, ENSBTAG00000039647, ENSBTAG00000052023, ENSBTAG00000053565, MGC152010 | 5.24E-10 | Protein percentage | ETH | DAM | Liver |
| CSN2 | 2.15E-07 | Protein percentage | INRAE | NOR | Mammary Gland |
| LAPTM4A | 3.23E-07 | Stature | USDA | HOL | Mammary Gland |
| PKLR | 6.36E-07 | Protein percentage | INRAE | HOL | Liver |
| TMPRSS11BNL | 1.62E-06 | Protein percentage | INRAE | NOR | Rumen |
| PKD2 | 1.57E-05 | Protein percentage | USDA | HOL | Mammary Gland |
| ABO | 4.69E-05 | Protein percentage | USDA | HOL | Jejunum |
| ENSBTAG00000049117, PUM2 | 5.98E-05 | Stature | USDA | HOL | Jejunum |
| CSF2RB | 0.000111005 | Protein percentage | ETH | DAM | Mammary Gland |
| ENSBTAG00000052400 | 0.00013592 | Protein percentage | INRAE | NOR | Mammary Gland |
| SELENOP, bta-mir-12004 | 0.000167438 | Protein percentage | USDA | HOL | Rumen |
| SLC52A2 | 0.00024349 | Milk yield | INRAE | HOL | Mammary Gland |
| CYHR1 | 0.000311285 | Milk yield | INRAE | HOL | Liver |
| MAFA | 0.000340009 | Protein percentage | INRAE | NOR | Muscle |
| ENSBTAG00000049824 | 0.000376683 | Protein percentage | INRAE | MON | Mammary Gland |
| LY6E | 0.000407975 | Milk yield | INRAE | HOL | Jejunum |
| SHE | 0.000411819 | Protein percentage | INRAE | HOL | Liver |
| RIN2 | 0.000496454 | Milk yield | USDA | HOL | Mammary Gland |
| LMAN1 | 0.001043752 | Protein percentage | INRAE | HOL | Mammary Gland |
| ENSBTAG00000003523 | 0.001104402 | Protein percentage | ETH | DAM | Liver |
| TTC38 | 0.001884925 | Protein percentage | INRAE | MON | Liver |
| Novel(3), PABPC1 | 0.00404533 | Protein percentage | INRAE | HOL | Rumen |
| ENSBTAG00000002148, TMEM205 | 0.004191098 | Stature | USDA | HOL | Mammary Gland |
| EED, Novel(1) | 0.006027843 | Protein percentage | INRAE | HOL | Mammary Gland |
| HMGCS1 | 0.006852071 | Protein percentage | USDA | HOL | Rumen |
| CHID1 | 0.007645734 | Stature | USDA | HOL | Mammary Gland |
| GRN | 0.008698239 | Protein percentage | INRAE | MON | Mammary Gland |
| GPIHBP1 | 0.008710602 | Protein percentage | USDA | HOL | Mammary Gland |
| ADAM15, EFNA4 | 0.016032205 | Protein percentage | INRAE | NOR | Rumen |
| CSN1S1, CSN1S2, CSN3 | 0.016485658 | Protein percentage | INRAE | MON | Mammary Gland |
| TALDO1 | 0.017416243 | Stature | USDA | HOL | Liver |
| PIGC | 0.027834882 | Protein percentage | INRAE | MON | Rumen |
| RPL29 | 0.034671816 | Protein percentage | INRAE | NOR | Rumen |
| PUF60 | 0.037273866 | Milk yield | USDA | HOL | Liver |
| UTP3 | 0.041959163 | Milk yield | ETH | DAM | Liver |
| Novel(17) | 0.043125269 | Protein percentage | USDA | HOL | Liver |
| PBXIP1 | 0.045780072 | Protein percentage | INRAE | NOR | Muscle |

Other genes with significant gene-trait associations with milk production in our analysis (Table 3) included SLC4A4, which was associated with expression in jejunum in this study, and has previously been linked to milk production and composition traits in Barka cattle [43]. Another gene associated with expression in the jejunum and linked to protein content, protein percentage and milk yield traits was ABO. ABO has been associated with milk production traits using GWAS in Holstein-Friesian cattle previously by Jiang et al. (2019) [28]. Jiang et al. (2019) also linked CSN1S1 to fat and protein percentage in milk which we found was significantly associated with these traits and expression in mammary gland in our analysis. The predicted expression of gene PKD2 in mammary gland tissue was associated with protein percentage, and this gene has previously been associated with milk production traits in Norwegian Red cattle [44]. CSF2RB also showed a significant gene-trait association with protein content and protein percentage and is known to be highly expressed in the mammary gland of lactating dairy cattle [45] and associated with fat and protein percentage [46].

Figure 10 shows the results for milk protein percentage using the lactating mammary gland PredictDB models. As can be seen for most GWAS loci a candidate driver gene is identified. Notably PICALM was significantly associated with milk protein percentage despite no significant association being observed at the same locus in the underlying GWAS data. Given PICALM has previously been linked to milk composition traits [47] this suggests that this TWAS result is likely not a spurious association. This highlights how TWAS can not only identify the genes potentially driving significant GWAS loci, but also identify new potential loci linked to the trait where no individual variant was genome-wide significant in the original GWAS data.

**Figure 10:**
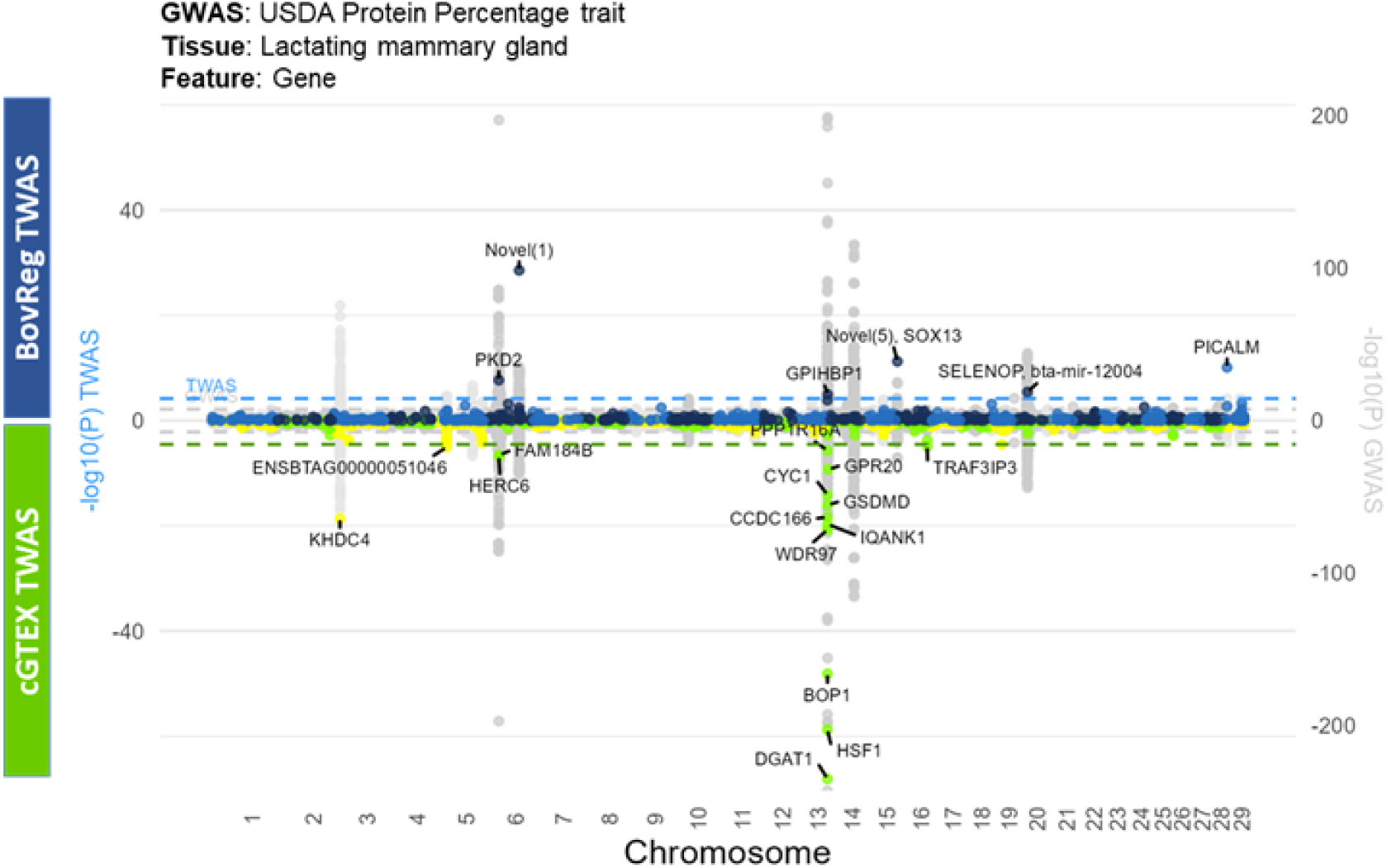
Comparison of CattleGTEx and BovReg TWAS results for the protein percentage trait in the mammary gland. The grey points represent USDA protein percentage GWAS results, with each point corresponding to a variant’s -log10(p-value), plotted by chromosomal position. Alternating background colors distinguish chromosomes. BovReg TWAS results (blue points) are shown in the upper half, while CattleGTEx TWAS results (yellow and green points) are in the lower half. This highlights overlapping and unique gene associations across datasets.

### Comparison of our TWAS results with results from the CattleGTEx project

We compared the results of our TWAS to those from the CattleGTEx project [19] (Figure 10). Despite overlaps these results are not redundant, with some GWAS peaks only being linked to a candidate gene in one or other dataset (Figure 10). For example, the KHDC4 locus, which has previously been associated with milk casein content in Belgian Blue cattle [48], was only identified in the CattleGTEx dataset, while, a novel gene on chromosome 6 and PICALM (mentioned above) were only identified in our analysis.

The CattleGTEx project identified a TWAS peak linked to DGAT1 [19]. DGAT1 is a well-known functional gene for fat content and milk production in cattle, though the main effect of the DGAT1 variants are not on protein synthesis and gene expression level [42,49]. In Figure 10 depicting the TWAS results for milk protein percentage and gene expression in the mammary gland tissue, a TWAS peak for DGAT1 is missing in the BovReg results because no valid model was generated for this gene in this tissue. Closer examination highlighted this is likely in part because the gene models for DGAT1 and SCRT are merged in the BovReg transcriptome annotation, whereas the CattleGTEx project used the corresponding Ensembl annotation. However, Figures 9 shows that a significant gene-trait association of this merged model with milk protein percentage at the gene level is still detected in the rumen. Together these results illustrate our results are complementary to the previous CattleGTEx results, with several new associations detected both at known and novel genes providing new insights into the biological mechanisms driving these phenotypes.

We then performed a more detailed comparative analysis, using mammary gland tissue as an example, of significant genes identified in our BovReg TWAS approach, and those in the CattleGTEx dataset within a 5 Megabase (Mb) window. With the aim of identifying how the different approaches and annotations used impact results. We observed several novel gene associations across various phenotypic categories in the BovReg cohort (Supplementary Table 11). These are described below for three subsets of traits:

### Milk Production and Composition Traits

For traits related to milk production and composition— namely milk yield, protein content, protein percentage, and fat percentage—we identified multiple significant genes unique to the BovReg dataset. Notably, MSTRG.7781 (RIN2) was significantly associated with both milk yield and protein content, suggesting a potential role in lactation processes. In the case of protein percentage, four out of six significant genes (66.67%) were unique, including MSTRG.38332 (Novel (1)), MSTRG.12002 (Novel (5), SOX13), MSTRG.29965 (PICALM), and MSTRG.21200 (SELENOP, bta-mir-12004). For fat percentage, MSTRG.13074 (IVNS1ABP) was the unique significant gene identified, potentially indicating a novel involvement in lipid metabolism within bovine mammary tissue.

### Body Conformation Traits

Regarding body conformation traits such as body depth and stature, we found that MSTRG.39515 (ENSBTAG00000002148, TMEM205) was uniquely associated with both traits in the BovReg dataset. Additionally, for stature, MSTRG.30950 (CHID1) and MSTRG.30962 (TALDO1) were identified as significant genes without corresponding significant CattleGTEx associated genes within 5 Mb, suggesting novel genetic influences on skeletal development in cattle.

### Reproductive Traits

In the category of reproductive traits, specifically sire stillbirth, the gene MSTRG.35987 (CNPY2, CS, bta-mir-12054) was uniquely significant in our dataset. This finding may point to new genetic factors affecting reproductive outcomes in bovines. Especially as CNPY2 was identified previously in a genome-wide scan for candidate genes underlying maternal effects of preweaning calf mortality in Nellore cattle [50].

These results indicate that due to the difference in annotations and RNA-seq and GWAS datasets used, a substantial proportion of significant gene associations identified in the BovReg TWAS using mammary gland tissue as an example are unique to our bovine cohort and were not detected in the CattleGTEx dataset within the specified genomic window. The recurrence of genes like MSTRG.7781 (RIN2) and MSTRG.39515 (TMEM205) across multiple related traits underscores their potential importance in bovine physiology and development.

## Discussion

We have provided gene-trait associations for cattle using a novel, reusable TWAS workflow that generates gene expression prediction models from RNA-seq datasets with or without matching genotype data and imputes gene expression levels into GWAS cohorts with just summary statistics. Since reference panels of matching gene expression and genotypes across large cohorts of cattle are rare and expensive to generate, as part of our workflow we included the ability to produce prediction models where the genotypes are called from the RNA-seq data directly. We show that genotypes called from RNA-seq data with software designed for low pass sequencing data are more accurate than those called using previously adopted methods. In the analysis presented in this study we tested the workflow we developed by training transcriptome models on genotypes called from publicly available large RNA-seq cohorts [6,39] and performed a TWAS using GWAS summary statistics provided by BovReg partners, and publicly available datasets [28].

The most significant gene-trait associations in our analysis were for milk yield and protein percentage and content. Reassuringly, several known gene-trait associations were identified including for DGAT1 [42,49], PKD2 [44], SLC4A4 [43], ABO [28], and CSN1S1 [28]. There were also some interesting novel associations for other traits identified. For example, the immunity related gene CD14 expressed in mammary gland tissue was associated with gestation length (GL-DYD). CD14 is essential for pathogen recognition in the mammary gland and other tissues and is expressed at high levels in bovine mammary epithelial cells and in milk [51]. In contrast for somatic cell count (SCS), the only immune related trait in our analysis, there were no gene-trait associations for genes with known immune function, though there were significant associations with genes with routine cellular processes including GPAA1 and CPSF1 expressed in liver and jejunum respectively. The only other gene with known immune function and a significant gene-trait association in our analysis was CSF2RB which was associated with protein content and milk yield in the mammary gland. Previous studies have shown that multiple QTL underlie milk phenotypes at the CSF2RB locus [46,52] and that CSF2RB is highly expressed in the mammary gland in dairy cattle [45]. The lack of immunity related gene-trait associations in our study is largely due to the traits analysed. Other recent studies have used TWAS to fine-map gene-trait associations for resistance to bovine tuberculosis [21] and to understand susceptibility to metabolic diseases such as ketosis [22].

We also highlight that several novel genes were discovered in our analyses that were undetected or untested in previous studies, including, when comparing our results to TWAS results from the CattleGTEx consortium [19], and even when using the same GWAS summary statistics. For example, the PICALM gene was linked to protein percentage in mammary gland tissue in our analysis but not in CattleGTEx, suggesting a difference in the gene expression prediction models between the two studies. Whereas CattleGTEx combined RNA-seq datasets deriving from different studies, in this analysis we chose to only use single cohort RNA-seq datasets. This has the disadvantage of reducing sample sizes but potentially reduces confounding by batch effects. Likewise, our use of a novel gene build (the BovReg transcriptome annotation) enabled us to detect associations with previously untested genes and transcripts, providing new insights into potentially causative genes. Notably, however, CattleGTEx did detect associations not identified in this study. Consequently, a combination of both approaches may produce the most complete set of genes and loci linked to given traits.

Due to a lack of available GWAS data, and/or summary statistics our analysis was focused predominantly on milk production traits. Sharing of summary statistics from GWAS would greatly facilitate TWAS studies for cattle and other farmed animals. Recent publications have described the potential benefits of sharing summary statistics [45] with suitable defined metadata standards [54]. The analysis frameworks we have provided, however, can now be expanded to other tissue types and traits e.g. for feed efficiency/conversion [16,55]. We have shared the Nextflow [56] workflows via GitHub (https://github.com/evotools/bovreg-twas-suite). Standardised analysis pipelines are an important part of FAIR bioinformatics research [57]. Recently a case study was published of nf-core adoption by the BovReg Project and five other European research consortia under the EuroFAANG umbrella and dedicated to farmed animal genomics [57]. Researchers can now apply our pipeline to their own datasets and assess their ability to identify novel gene-trait associations using these models and appropriate summary statistics across mapping populations. The gene expression models we have generated have been shared with partners within the BovReg consortium to assess their merit in predicting phenotypes and genomic breeding values and to identify loci of interest for functional validation in appropriate bovine cell lines.

The functional annotation information generated for cattle, by the BovReg project e.g. [29], US Cattle FAANG projects e.g. [58] and shared via the Functional Annotation of Animal Genomes (FAANG) initiative provides the opportunity to understand transcriptional/post-transcriptional regulatory mechanisms underpinning GWAS and TWAS hits for complex traits in cattle [59,60]. To facilitate further analysis gene-trait associations could be made available as tracks, through the FAANG Data Portal (https://data.faang.org/home) [61] and the Ensembl genome browser [62], including, for example, the chromosomal locations, positions and lead SNPs. Our comprehensive Nextflow pipeline offers a reusable, reproducible, and transparent solution for transcriptome-wide association studies across species—a resource that will fuel ongoing and future FAANG efforts aimed at linking functional annotation to phenotypes. By combining updated genome annotations with robust gene expression prediction models, we provide insights into the regulatory architecture of complex traits in cattle, a priority species for the FAANG community. We share the pipeline and resources openly, dovetailing with FAANG’s commitment to FAIR (Findable, Accessible, Interoperable, and Reusable) data and tool development. The results and the workflow we have presented for the BovReg project will contribute to improving our understanding of the molecular drivers of complex traits in cattle populations with the goal of eventually leveraging this information in breeding decisions.

## Methods

### Reference Panel Generation for Genotype Imputation

The reference panel used for genotype imputation from RNA-seq data was generated from publicly available genotypes provided by the 1000 Bull Genomes Project (accession number ERZ1738264). To ensure compatibility with GLIMPSE and to maintain data quality, the genotype data were initially filtered for biallelic SNPs located on autosomes using *bcftools* (v1.13). Further filtering steps included applying call rate (CR) and genotype quality (GQ) thresholds with *bcftools*, retaining only SNPs with CR ≥ 75% and GQ ≥ 20. Finally, the filtered reference panel was phased using BEAGLE [31].

### Evaluation of Imputation Accuracy

The imputation accuracy of GLIMPSE v1.1.1 [32] was evaluated in comparison to BEAGLE using matching whole-genome sequencing (WGS) genotype calls and RNA-seq data from [33]. The WGS genotypes included three Holstein (European Taurine: HF3457, HF3458, HF3471) and three Nellore (Indicine: Nellore1, Nellore2, Nellore3) samples. RNA was derived from various immune cell types (B cells, CD4, CD8, Monocytes, NK cells) and deeply sequenced, yielding up to 600 million paired-end RNA-seq reads per individual.

To assess imputation accuracy across different sequencing depths, reflective of those commonly found in public datasets, the merged RNA-seq fastq reads across cell types were downsampled using Seqtk (v1.3 (r106)), maintaining pairing consistency with a fixed random seed. The reads were downsampled to 15 million, 30 million, and 60 million reads, and the results were compared to both the WGS genotypes and imputed calls.

### RNA-seq Data Processing and Genotype Calling

RNA-seq reads were aligned to the ARS-UCD1.2 reference genome using the Ensembl v106 gene build for GLIMPSE validation and the BovReg transcriptome annotation for downstream processing of study samples with STAR (version 2.7.10a) [25], applying specific parameters for alignment (outFilterMismatchNmax 3, outFilterMultimapNmax 10, outFilterScoreMinOverLread 0.66). Initial genotypes were called using *bcftools* (v1.13) mpileup. Imputation was performed using the filtered reference panel with GLIMPSE v1.1.1 and BEAGLE. Concordance was then calculated for homozygous reference alleles (0/0) and non-reference alleles (0/1, 1/1) after filtering the genotypes for WGS GQ > 25 and imputed genotypes with GP ≥ 0.8.

### RNA-seq Data Processing Pipeline

A Nextflow pipeline was implemented to process RNA-seq reads, quantify gene and transcript expression levels, and impute genotypes for training PredictDB models. The pipeline accepts input in the form of publicly available RNA-seq accession IDs or local fastq files, along with a reference genome, gene annotation, and a genotype reference panel.

The process begins by downloading the fastq files associated with the accession IDs using enaBrowserTools (github.com/enasequence/enaBrowserTools). These files undergo quality control and adapter sequence trimming via TrimGalore (v 0.6.7) [63]. The trimmed reads are then aligned to the reference genome using STAR (v2.7.10a, parameters: outFilterMismatchNmax 3, outFilterMultimapNmax 10, outFilterScoreMinOverLread 0.66) for gene-level quantification, while transcript-level quantification is performed using Kallisto (v 0.50.0, option:-b 100) [35]. Genes and transcripts are filtered to include only those expressed in at least 50% of the samples. Covariates are estimated from the expression data using PEER tools (r-peer v1.3) [36], with the number of PEER factors determined based on sample size: 60 factors for ≥350 samples, 45 for 250–349 samples, 30 for 150–249 samples, and 15 for fewer than 150 samples.

### PredictDB Model Training

The training process utilized a custom R script adapted from the PredictDB repository (github.com/hakyimlab/PredictDB-Tutorial). This script employs the glmnet library for elastic net regularization, processing genotype and expression data by filtering SNPs, adjusting gene expression for covariates, and conducting cross-validated elastic net regression. Only SNPs that overlapped between the genotype data and those tested in the GWAS study were included. Importantly, the set of included SNPs varied across cohorts due to differences in genotype quality control, and availability in the GWAS datasets. This resulted in cohort-specific variant sets being used for model training, tailored to maximize performance within each respective cohort. Model performance was evaluated using R-squared, correlation, and p-values, and models were filtered based on performance criteria (z-score p-value < 0.05 and rho average > 0.1).

### Transcriptome-wide association studies

The resulting PredictDB models were subsequently used with S-PrediXcan (MetaXcan 0.7.5) [41] to analyze TWAS results using available GWAS summary statistics. TWAS result p-values were corrected using the Bonferroni adjustment procedure, and significant hits were identified for each GWAS study cohort.

### Annotation of BovReg transcript model using Ensembl Reference Annotations

To integrate the BovReg RNA-Seq transcriptome with Ensembl reference annotations (Bos taurus ARS-UCD1.2, release 111), we used gffcompare v0.12.6 to classify assembled transcripts based on their alignment with known genes. Transcripts were systematically annotated with gene IDs, gene names, and reference transcript mappings. Novel transcripts were assigned unique labels, with the number in parentheses indicating how many novel transcripts were identified for each gene. The final annotation file provides a comprehensive mapping of BovReg transcripts to reference genes, distinguishing known and novel transcriptomic features in cattle.

## Author contributions

SJ developed the TWAS analysis workflow, analysed the data, produced most of the figures, and wrote the initial draft of the manuscript. ELC and JP designed the study and co-wrote the manuscript with SJ. JP also contributed to generation of the figures. GCMM and CC prepared the gtf annotation file from the transcriptomic data they generated for the BovReg Project. MS contributed to generating the gtf annotation file. CK, HP, DB, and M-PS provided summary statistics for the GWAS cohorts analysed in the study. NKK provided advice on the TWAS workflow. PKC ran the TWAS workflow on the GWAS and matched RNA-Seq datasets provided by FBN. CK coordinated the BovReg project. All co-authors provided feedback on the initial draft of the manuscript and read and approved the final version of the manuscript before submission.

## Code availability

The Nextflow workflows are available via GitHub (https://github.com/evotools/bovreg-twas-suite).

## Data Availability

To test the TWAS workflow we used publicly available datasets, which are described, including accession numbers, within the manuscript above.

The supplementary figures are available via https://figshare.com/articles/figure/BovReg_TWAS_Supplementary_Figures_for_manuscript_A_novel_reusable_transcriptome-wide_association_study_workflow_used_to_map_key_genes_linked_to_important_cattle_traits/29264795?file=55207907 and the supplementary tables via https://figshare.com/articles/dataset/BovReg_TWAS_Supplementary_Tables_for_manuscript_A_novel_reusable_transcriptome-wide_association_study_workflow_used_to_map_key_genes_linked_to_important_cattle_traits/29264792?file=55207901.

The new transcriptome annotation file, used for this analysis generated by BovReg partners (BovReg_RNA-Seq_CAGE_merged_track_sorted_LIEGE.gtf) is available via https://figshare.com/articles/dataset/BovReg_TWAS_Annotation_File_BovReg_RNA-Seq_CAGE_merged_track_sorted_LIEGE_gtf/29264816?file=55229522.

The summary statistics from GWAS provided by the BovReg partners are available upon request.

## Supplementary Materials

Supplementary Table 1: GWAS sources, phenotypes, cohort sizes, and breed information used in the study.

Supplementary Table 2: Details of the RNA-seq datasets used for model training. It includes accession IDs, tissue types, SNP sets, and the number of models pre- and post-filtering based on z-score p-value (<0.05) and rho average (>0.1).

Supplementary Table 3: BovReg Gene Models along with their corresponding GffCompare hits. It includes comparisons to known gene annotations.

Supplementary Table 4: BovReg IDs and their best matching transcript annotations. It identifies the most accurate matches for each transcript model, including novel and known annotations.

Supplementary Table 5: These set of tables contain the summary of significant gene and transcript associations across partner-breed combinations and tissues for each TWAS.

Supplementary Table 6: The top genes associated with milk yield across more than one tissue or cohort.

Supplementary Table 7: The top genes associated with protein percentage across more than one tissue or cohort.

Supplementary Table 8: The top genes associated with stature across more than one tissue or cohort. Supplementary Table 9: Top hit genes with Bonferroni-adjusted p-value < 0.05 across all TWAS. Supplementary Table 10: Top hit transcripts with Bonferroni-adjusted p-value < 0.05 across all TWAS.

Supplementary Table 11: Comparison of novel TWAS hits between BovReg and CattleGTEx in the mammary gland across phenotypes.

Supplementary Figure 1: GLIMPSE imputation performed comparatively poorly on the Bos indicus samples, in particular for rarer variants.

Supplementary Figure 2: ETH Gene model counts shared across tissues for PredictDB trained models (zscore pval <0.05 and rho avg.>0.1).

Supplementary Figure 3: ETH transcript model counts shared across tissues for PredictDB trained models (zscore pval <0.05 and rho avg.>0.1)

Supplementary Figure 4: INRAE Gene model counts shared across tissues for PredictDB trained models (zscore pval <0.05 and rho avg.>0.1)

Supplementary Figure 5: INRAE transcript model counts shared across tissues for PredictDB trained models (zscore pval <0.05 and rho avg.>0.1)

Supplementary Figure 6: FBN gene model counts shared across tissues for PredictDB trained models (zscore pval <0.05 and rho avg.>0.1)

Supplementary Figure 7: FBN transcript model counts shared across tissues for PredictDB trained models (zscore pval <0.05 and rho avg.>0.1)

Supplementary Figure 8: Visualisation of genes with significant TWAS associations across each of the five tissues types for all of the traits analysed (p-value <0.05, Bonferroni corrected). Integration of the SNP genotyping, gene expression, and GWAS summary data using S-PrediXcan. White areas indicate no valid model was trained for the gene indicated on the X axis in the given tissue (labelled to the right). Grey areas mean a valid model was trained but the gene’s expression was not significantly associated with the trait indicated on the left Y axis. Coloured boxes indicate the gene’s expression was significantly associated with the given trait in the specified tissue.

Supplementary Figure 9: Visualisation of transcripts with significant TWAS associations across each of the five tissues analysed for the traits analysed (BH corrected p-value <0.05). Integration the SNP genotyping, transcript expression, and GWAS summary data using S-PrediXcan.

Supplementary Figure 10: Transcripts with significant TWAS associations summarised for each of the traits analysed (BH corrected p-value <0.05). Overlaps in tissue-specificity, chromosomal location and start position are shown for each transcript.

## References

[1] G. Sahana, Z. Cai, M.P. Sanchez, A.C. Bouwman, D. Boichard, Invited review: Good practices in genome-wide association studies to identify candidate sequence variants in dairy cattle, Journal of Dairy Science 106 (2023) 5218–5241. 10.3168/jds.2022-22694.

[2] M.E. Goddard, B.J. Hayes, Genome-Wide Association Studies and Linkage Disequilibrium in Cattle, in: Bovine Genomics, 2012: pp. 192–210. 10.1002/9781118301739.ch13.

[3] J.I. Weller, Mapping Quantitative Trait Loci, in: Bovine Genomics, 2012: pp. 169–191. 10.1002/9781118301739.ch12.

[4] Z.-L. Hu, C.A. Park, J.M. Reecy, Building a livestock genetic and genomic information knowledgebase through integrative developments of Animal QTLdb and CorrDB, Nucleic Acids Research 47 (2019) D701–D710. 10.1093/nar/gky1084.

[5] A.C. Nica, E.T. Dermitzakis, Expression quantitative trait loci: present and future., Philos Trans R Soc Lond B Biol Sci 368 (2013) 20120362. 10.1098/rstb.2012.0362.

[6] T.J. Lopdell, K. Tiplady, M. Struchalin, T.J.J. Johnson, M. Keehan, R. Sherlock, C. Couldrey, S.R. Davis, R.G. Snell, R.J. Spelman, M.D. Littlejohn, DNA and RNA-sequence based GWAS highlights membrane-transport genes as key modulators of milk lactose content, BMC Genomics 18 (2017) 968. 10.1186/s12864-017-4320-3.

[7] Y. Tang, J. Zhang, W. Li, X. Liu, S. Chen, S. Mi, J. Yang, J. Teng, L. Fang, Y. Yu, Identification and characterization of whole blood gene expression and splicing quantitative trait loci during early to mid-lactation of dairy cattle, BMC Genomics 25 (2024) 445. 10.1186/s12864-024-10346-7.

[8] M.D. Littlejohn, K. Tiplady, T.A. Fink, K. Lehnert, T. Lopdell, T. Johnson, C. Couldrey, M. Keehan, R.G. Sherlock, C. Harland, A. Scott, R.G. Snell, S.R. Davis, R.J. Spelman, Sequence-based Association Analysis Reveals an MGST1 eQTL with Pleiotropic Effects on Bovine Milk Composition, Scientific Reports 6 (2016) 25376. 10.1038/srep25376.

[9] I. van den Berg, B.J. Hayes, A.J. Chamberlain, M.E. Goddard, Overlap between eQTL and QTL associated with production traits and fertility in dairy cattle, BMC Genomics 20 (2019) 291. 10.1186/s12864-019-5656-7.

[10] W. Dong, J. Yang, Y. Zhang, S. Liu, C. Ning, X. Ding, W. Wang, Y. Zhang, Q. Zhang, L. Jiang, Integrative analysis of genome-wide DNA methylation and gene expression profiles reveals important epigenetic genes related to milk production traits in dairy cattle, Journal of Animal Breeding and Genetics 138 (2021) 562–573. 10.1111/jbg.12530.

[11] M. Forutan, B.N. Engle, A.J. Chamberlain, E.M. Ross, L.T. Nguyen, M.J. D’Occhio, A.C. Snr, E.A. Kho, G. Fordyce, S. Speight, M.E. Goddard, B.J. Hayes, Genome-wide association and expression quantitative trait loci in cattle reveals common genes regulating mammalian fertility, Communications Biology 7 (2024) 724. 10.1038/s42003-024-06403-2.

[12] W. Cai, Y. Zhang, T. Chang, Z. Wang, B. Zhu, Y. Chen, X. Gao, L. Xu, L. Zhang, H. Gao, J. Song, J. Li, The eQTL colocalization and transcriptome-wide association study identify potentially causal genes responsible for economic traits in Simmental beef cattle, Journal of Animal Science and Biotechnology 14 (2023) 78. 10.1186/s40104-023-00876-7.

[13] G.M. Guillocheau, A. El Hou, C. Meersseman, D. Esquerré, E. Rebours, R. Letaief, M. Simao, N. Hypolite, E. Bourneuf, N. Bruneau, A. Vaiman, C.J. Vander Jagt, A.J. Chamberlain, D. Rocha, Survey of allele specific expression in bovine muscle, Scientific Reports 9 (2019) 4297. 10.1038/s41598-019-40781-6.

[14] I.S. Garcia, B. Silva-Vignato, A.S.M. Cesar, J. Petrini, V.H. da Silva, N.S. Morosini, C.P. Goes, J. Afonso, T.R. da Silva, B.D. Lima, L.G. Clemente, L.C. de A. Regitano, G.B. Mourão, L.L. Coutinho, Novel putative causal mutations associated with fat traits in Nellore cattle uncovered by eQTLs located in open chromatin regions, Scientific Reports 14 (2024) 10094. 10.1038/s41598-024-60703-5.

[15] J.J. Bruscadin, M.M. de Souza, K.S. de Oliveira, M.I.P. Rocha, J. Afonso, T.F. Cardoso, A. Zerlotini, L.L. Coutinho, S.C.M. Niciura, L.C. de Almeida Regitano, Muscle allele-specific expression QTLs may affect meat quality traits in Bos indicus, Scientific Reports 11 (2021) 7321. 10.1038/s41598-021-86782-2.

[16] C. McKenna, K. Keogh, R.K. Porter, S.M. Waters, P. Cormican, D.A. Kenny, An examination of skeletal muscle and hepatic tissue transcriptomes from beef cattle divergent for residual feed intake, Scientific Reports 11 (2021) 8942. 10.1038/s41598-021-87842-3.

[17] J. Mai, M. Lu, Q. Gao, J. Zeng, J. Xiao, Transcriptome-wide association studies: recent advances in methods, applications and available databases, Communications Biology 6 (2023) 899. 10.1038/s42003-023-05279-y.

[18] Y. Hu, M. Li, Q. Lu, H. Weng, J. Wang, S.M. Zekavat, Z. Yu, B. Li, J. Gu, S. Muchnik, Y. Shi, B.W. Kunkle, S. Mukherjee, P. Natarajan, A. Naj, A. Kuzma, Y. Zhao, P.K. Crane, H. Lu, H. Zhao, Alzheimer’s Disease Genetics Consortium, A statistical framework for cross-tissue transcriptome-wide association analysis, Nature Genetics 51 (2019) 568–576. 10.1038/s41588-019-0345-7.

[19] S. Liu, Y. Gao, O. Canela-Xandri, S. Wang, Y. Yu, W. Cai, B. Li, R. Xiang, A.J. Chamberlain, E. Pairo-Castineira, K. D’Mellow, K. Rawlik, C. Xia, Y. Yao, P. Navarro, D. Rocha, X. Li, Z. Yan, C. Li, B.D. Rosen, C.P. Van Tassell, P.M. Vanraden, S. Zhang, L. Ma, J.B. Cole, G.E. Liu, A. Tenesa, L. Fang, A multi-tissue atlas of regulatory variants in cattle, Nature Genetics 54 (2022) 1438–1447. 10.1038/s41588-022-01153-5.

[20] X.M. Mapel, N.K. Kadri, A.S. Leonard, Q. He, A. Lloret-Villas, M. Bhati, M. Hiltpold, H. Pausch, Molecular quantitative trait loci in reproductive tissues impact male fertility in cattle, Nature Communications 15 (2024) 674. 10.1038/s41467-024-44935-7.

[21] J.F. O’Grady, G.P. McHugo, J.A. Ward, T.J. Hall, S.L. Faherty O’Donnell, C.N. Correia, J.A. Browne, M. McDonald, E. Gormley, V. Riggio, J.G.D. Prendergast, E.L. Clark, H. Pausch, K.G. Meade, I.C. Gormley, S.V. Gordon, D.E. MacHugh, Integrative genomics sheds light on the immunobiology of tuberculosis in cattle, bioRxiv (2024) 2024.02.27.582295. 10.1101/2024.02.27.582295.

[22] Z. Yan, H. Huang, E. Freebern, D.J.A. Santos, D. Dai, J. Si, C. Ma, J. Cao, G. Guo, G.E. Liu, L. Ma, L. Fang, Y. Zhang, Integrating RNA-Seq with GWAS reveals novel insights into the molecular mechanism underpinning ketosis in cattle, BMC Genomics 21 (2020) 489. 10.1186/s12864-020-06909-z.

[23] A.S. Leonard, X.M. Mapel, H. Pausch, RNA sequencing variants are enriched for eQTL in cattle tissues, bioRxiv (2024) 2024.04.29.591607. 10.1101/2024.04.29.591607.

[24] S. Rubinacci, D.M. Ribeiro, R.J. Hofmeister, O. Delaneau, Efficient phasing and imputation of low-coverage sequencing data using large reference panels, Nature Genetics 53 (2021) 120–126. 10.1038/s41588-020-00756-0.

[25] A. Dobin, C.A. Davis, F. Schlesinger, J. Drenkow, C. Zaleski, S. Jha, P. Batut, M. Chaisson, T.R. Gingeras, STAR: ultrafast universal RNA-seq aligner., Bioinformatics 29 (2013) 15–21. 10.1093/bioinformatics/bts635.

[26] E.R. Gamazon, H.E. Wheeler, K.P. Shah, S.V. Mozaffari, K. Aquino-Michaels, R.J. Carroll, A.E. Eyler, J.C. Denny, D.L. Nicolae, N.J. Cox, H.K. Im, GTEx Consortium, A gene-based association method for mapping traits using reference transcriptome data, Nature Genetics 47 (2015) 1091–1098. 10.1038/ng.3367.

[27] A.N. Barbeira, M. Pividori, J. Zheng, H.E. Wheeler, D.L. Nicolae, H.K. Im, Integrating predicted transcriptome from multiple tissues improves association detection., PLoS Genet 15 (2019) e1007889. 10.1371/journal.pgen.1007889.

[28] J. Jiang, J.B. Cole, E. Freebern, Y. Da, P.M. VanRaden, L. Ma, Functional annotation and Bayesian fine-mapping reveals candidate genes for important agronomic traits in Holstein bulls, Communications Biology 2 (2019) 212. 10.1038/s42003-019-0454-y.

[29] M. Salavati, R. Clark, D. Becker, C. Kühn, G. Plastow, S. Dupont, G.C.M. Moreira, C. Charlier, E.L. Clark, Improving the annotation of the cattle genome by annotating transcription start sites in a diverse set of tissues and populations using Cap Analysis Gene Expression sequencing., G3 (Bethesda) 13 (2023). 10.1093/g3journal/jkad108.

[30] R. Poplin, V. Ruano-Rubio, M.A. DePristo, T.J. Fennell, M.O. Carneiro, G.A. Van der Auwera, D.E. Kling, L.D. Gauthier, A. Levy-Moonshine, D. Roazen, K. Shakir, J. Thibault, S. Chandran, C. Whelan, M. Lek, S. Gabriel, M.J. Daly, B. Neale, D.G. MacArthur, E. Banks, Scaling accurate genetic variant discovery to tens of thousands of samples, bioRxiv (2018) 201178. 10.1101/201178.

[31] B.L. Browning, X. Tian, Y. Zhou, S.R. Browning, Fast two-stage phasing of large-scale sequence data, The American Journal of Human Genetics 108 (2021) 1880–1890. 10.1016/j.ajhg.2021.08.005.

[32] S. Rubinacci, D.M. Ribeiro, R.J. Hofmeister, O. Delaneau, Efficient phasing and imputation of low-coverage sequencing data using large reference panels, Nature Genetics 53 (2021) 120–126. 10.1038/s41588-020-00756-0.

[33] J. Powell, A. Talenti, A. Fisch, J.D. Hemmink, E. Paxton, P. Toye, I. Santos, B.R. Ferreira, T.K. Connelley, L.J. Morrison, J.G.D. Prendergast, Profiling the immune epigenome across global cattle breeds, Genome Biology 24 (2023) 127. 10.1186/s13059-023-02964-3.

[34] B.J. Hayes, H.D. Daetwyler, 1000 Bull Genomes Project to Map Simple and Complex Genetic Traits in Cattle: Applications and Outcomes, Annual Review of Animal Biosciences 7 (2019) 89–102. 10.1146/annurev-animal-020518-115024.

[35] N.L. Bray, H. Pimentel, P. Melsted, L. Pachter, Near-optimal probabilistic RNA-seq quantification, Nature Biotechnology 34 (2016) 525–527. 10.1038/nbt.3519.

[36] O. Stegle, L. Parts, R. Durbin, J. Winn, A Bayesian framework to account for complex non-genetic factors in gene expression levels greatly increases power in eQTL studies., PLoS Comput Biol 6 (2010) e1000770. 10.1371/journal.pcbi.1000770.

[37] P. Danecek, J.K. Bonfield, J. Liddle, J. Marshall, V. Ohan, M.O. Pollard, A. Whitwham, T. Keane, S.A. McCarthy, R.M. Davies, H. Li, Twelve years of SAMtools and BCFtools, GigaScience 10 (2021) giab008. 10.1093/gigascience/giab008.

[38] C.P. Prowse-Wilkins, T.J. Lopdell, R. Xiang, C.J. Vander Jagt, M.D. Littlejohn, A.J. Chamberlain, M.E. Goddard, Genetic variation in histone modifications and gene expression identifies regulatory variants in the mammary gland of cattle, BMC Genomics 23 (2022) 815. 10.1186/s12864-022-09002-9.

[39] W. Nolte, R. Weikard, R.M. Brunner, E. Albrecht, H.M. Hammon, A. Reverter, C. Kühn, Biological Network Approach for the Identification of Regulatory Long Non-Coding RNAs Associated With Metabolic Efficiency in Cattle, Frontiers in Genetics 10 (2019). https://www.frontiersin.org/journals/genetics/articles/10.3389/fgene.2019.01130.

[40] F. Aguet, A.A. Brown, S.E. Castel, J.R. Davis, Y. He, B. Jo, P. Mohammadi, Y. Park, P. Parsana, A.V. Segrè, B.J. Strober, Z. Zappala, B.B. Cummings, E.T. Gelfand, K. Hadley, K.H. Huang, M. Lek, X. Li, J.L. Nedzel, D.Y. Nguyen, M.S. Noble, T.J. Sullivan, T. Tukiainen, D.G. MacArthur, G. Getz, A. Addington, P. Guan, S. Koester, A.R. Little, N.C. Lockhart, H.M. Moore, A. Rao, J.P. Struewing, S. Volpi, L.E. Brigham, R. Hasz, M. Hunter, C. Johns, M. Johnson, G. Kopen, W.F. Leinweber, J.T. Lonsdale, A. McDonald, B. Mestichelli, K. Myer, B. Roe, M. Salvatore, S. Shad, J.A. Thomas, G. Walters, M. Washington, J. Wheeler, J. Bridge, B.A. Foster, B.M. Gillard, E. Karasik, R. Kumar, M. Miklos, M.T. Moser, S.D. Jewell, R.G. Montroy, D.C. Rohrer, D. Valley, D.C. Mash, D.A. Davis, L. Sobin, M.E. Barcus, P.A. Branton, N.S. Abell, B. Balliu, O. Delaneau, L. Frésard, E.R. Gamazon, D. Garrido-Martín, A.D.H. Gewirtz, G. Gliner, M.J. Gloudemans, B. Han, A.Z. He, F. Hormozdiari, X. Li, B. Liu, E.Y. Kang, I.C. McDowell, H. Ongen, J.J. Palowitch, C.B. Peterson, G. Quon, S. Ripke, A. Saha, A.A. Shabalin, T.C. Shimko, J.H. Sul, N.A. Teran, E.K. Tsang, H. Zhang, Y.-H. Zhou, C.D. Bustamante, N.J. Cox, R. Guigó, M. Kellis, M.I. McCarthy, D.F. Conrad, E. Eskin, G. Li, A.B. Nobel, C. Sabatti, B.E. Stranger, X. Wen, F.A. Wright, K.G. Ardlie, E.T. Dermitzakis, T. Lappalainen, F. Aguet, K.G. Ardlie, B.B. Cummings, E.T. Gelfand, G. Getz, K. Hadley, R.E. Handsaker, K.H. Huang, S. Kashin, K.J. Karczewski, M. Lek, X. Li, D.G. MacArthur, J.L. Nedzel, D.T. Nguyen, M.S. Noble, A.V. Segrè, C.A. Trowbridge, T. Tukiainen, N.S. Abell, B. Balliu, R. Barshir, O. Basha, A. Battle, G.K. Bogu, A. Brown, C.D. Brown, S.E. Castel, L.S. Chen, C. Chiang, D.F. Conrad, N.J. Cox, F.N. Damani, J.R. Davis, O. Delaneau, E.T. Dermitzakis, B.E. Engelhardt, E. Eskin, P.G. Ferreira, L. Frésard, E.R. Gamazon, D. Garrido-Martín, A.D.H. Gewirtz, G. Gliner, M.J. Gloudemans, R. Guigo, I.M. Hall, B. Han, Y. He, F. Hormozdiari, C. Howald, H. Kyung Im, B. Jo, E. Yong Kang, Y. Kim, S. Kim-Hellmuth, T. Lappalainen, G. Li, X. Li, B. Liu, S. Mangul, M.I. McCarthy, I.C. McDowell, P. Mohammadi, J. Monlong, S.B. Montgomery, M. Muñoz-Aguirre, A.W. Ndungu, D.L. Nicolae, A.B. Nobel, M. Oliva, H. Ongen, J.J. Palowitch, N. Panousis, P. Papasaikas, Y. Park, P. Parsana, A.J. Payne, C.B. Peterson, J. Quan, F. Reverter, C. Sabatti, A. Saha, M. Sammeth, A.J. Scott, A.A. Shabalin, R. Sodaei, M. Stephens, B.E. Stranger, B.J. Strober, J.H. Sul, E.K. Tsang, S. Urbut, M. van de Bunt, G. Wang, X. Wen, F.A. Wright, H.S. Xi, E. Yeger-Lotem, Z. Zappala, J.B. Zaugg, Y.-H. Zhou, J.M. Akey, D. Bates, J. Chan, L.S. Chen, M. Claussnitzer, K. Demanelis, M. Diegel, J.A. Doherty, A.P. Feinberg, M.S. Fernando, J. Halow, K.D. Hansen, E. Haugen, P.F. Hickey, L. Hou, F. Jasmine, R. Jian, L. Jiang, A. Johnson, R. Kaul, M. Kellis, M.G. Kibriya, K. Lee, J. Billy Li, Q. Li, X. Li, J. Lin, S. Lin, S. Linder, C. Linke, Y. Liu, M.T. Maurano, B. Molinie, S.B. Montgomery, J. Nelson, F.J. Neri, M. Oliva, Y. Park, B.L. Pierce, N.J. Rinaldi, L.F. Rizzardi, R. Sandstrom, A. Skol, K.S. Smith, M.P. Snyder, J. Stamatoyannopoulos, B.E. Stranger, H. Tang, E.K. Tsang, L. Wang, M. Wang, N. Van Wittenberghe, F. Wu, R. Zhang, C.R. Nierras, P.A. Branton, L.J. Carithers, P. Guan, H.M. Moore, A. Rao, J.B. Vaught, S.E. Gould, N.C. Lockart, C. Martin, J.P. Struewing, S. Volpi, A.M. Addington, S.E. Koester, A.R. Little, Gte. Consortium, L. Analysts:, D.A.& C.C. (LDACC): Laboratory, N.I.H. program Management:, B. Collection:, Pathology:, eQTL manuscript working Group:, D.A.& C.C. (LDACC)— Analysis W.G. Laboratory, S.M. groups—Analysis W. Group, E.Gte. (eGTEx) Groups, N.I.H.C. Fund, NIH/NCI, NIH/NHGRI, NIH/NIMH, NIH/NIDA, B.C.S. Site—NDRI, Genetic effects on gene expression across human tissues, Nature 550 (2017) 204–213. 10.1038/nature24277.

[41] A.N. Barbeira, S.P. Dickinson, R. Bonazzola, J. Zheng, H.E. Wheeler, J.M. Torres, E.S. Torstenson, K.P. Shah, T. Garcia, T.L. Edwards, E.A. Stahl, L.M. Huckins, F. Aguet, K.G. Ardlie, B.B. Cummings, E.T. Gelfand, G. Getz, K. Hadley, R.E. Handsaker, K.H. Huang, S. Kashin, K.J. Karczewski, M. Lek, X. Li, D.G. MacArthur, J.L. Nedzel, D.T. Nguyen, M.S. Noble, A.V. Segrè, C.A. Trowbridge, T. Tukiainen, N.S. Abell, B. Balliu, R. Barshir, O. Basha, A. Battle, G.K. Bogu, A. Brown, C.D. Brown, S.E. Castel, L.S. Chen, C. Chiang, D.F. Conrad, F.N. Damani, J.R. Davis, O. Delaneau, E.T. Dermitzakis, B.E. Engelhardt, E. Eskin, P.G. Ferreira, L. Frésard, E.R. Gamazon, D. Garrido-Martín, A.D.H. Gewirtz, G. Gliner, M.J. Gloudemans, R. Guigo, I.M. Hall, B. Han, Y. He, F. Hormozdiari, C. Howald, B. Jo, E.Y. Kang, Y. Kim, S. Kim-Hellmuth, T. Lappalainen, G. Li, X. Li, B. Liu, S. Mangul, M.I. McCarthy, I.C. McDowell, P. Mohammadi, J. Monlong, S.B. Montgomery, M. Muñoz-Aguirre, A.W. Ndungu, A.B. Nobel, M. Oliva, H. Ongen, J.J. Palowitch, N. Panousis, P. Papasaikas, Y. Park, P. Parsana, A.J. Payne, C.B. Peterson, J. Quan, F. Reverter, C. Sabatti, A. Saha, M. Sammeth, A.J. Scott, A.A. Shabalin, R. Sodaei, M. Stephens, B.E. Stranger, B.J. Strober, J.H. Sul, E.K. Tsang, S. Urbut, M. van de Bunt, G. Wang, X. Wen, F.A. Wright, H.S. Xi, E. Yeger-Lotem, Z. Zappala, J.B. Zaugg, Y.-H. Zhou, J.M. Akey, D. Bates, J. Chan, L.S. Chen, M. Claussnitzer, K. Demanelis, M. Diegel, J.A. Doherty, A.P. Feinberg, M.S. Fernando, J. Halow, K.D. Hansen, E. Haugen, P.F. Hickey, L. Hou, F. Jasmine, R. Jian, L. Jiang, A. Johnson, R. Kaul, M. Kellis, M.G. Kibriya, K. Lee, J.B. Li, Q. Li, X. Li, J. Lin, S. Lin, S. Linder, C. Linke, Y. Liu, M.T. Maurano, B. Molinie, S.B. Montgomery, J. Nelson, F.J. Neri, M. Oliva, Y. Park, B.L. Pierce, N.J. Rinaldi, L.F. Rizzardi, R. Sandstrom, A. Skol, K.S. Smith, M.P. Snyder, J. Stamatoyannopoulos, B.E. Stranger, H. Tang, E.K. Tsang, L. Wang, M. Wang, N. Van Wittenberghe, F. Wu, R. Zhang, C.R. Nierras, P.A. Branton, L.J. Carithers, P. Guan, H.M. Moore, A. Rao, J.B. Vaught, S.E. Gould, N.C. Lockart, C. Martin, J.P. Struewing, S. Volpi, A.M. Addington, S.E. Koester, A.R. Little, L.E. Brigham, R. Hasz, M. Hunter, C. Johns, M. Johnson, G. Kopen, W.F. Leinweber, J.T. Lonsdale, A. McDonald, B. Mestichelli, K. Myer, B. Roe, M. Salvatore, S. Shad, J.A. Thomas, G. Walters, M. Washington, J. Wheeler, J. Bridge, B.A. Foster, B.M. Gillard, E. Karasik, R. Kumar, M. Miklos, M.T. Moser, S.D. Jewell, R.G. Montroy, D.C. Rohrer, D.R. Valley, D.A. Davis, D.C. Mash, A.H. Undale, A.M. Smith, D.E. Tabor, N.V. Roche, J.A. McLean, N. Vatanian, K.L. Robinson, L. Sobin, M.E. Barcus, K.M. Valentino, L. Qi, S. Hunter, P. Hariharan, S. Singh, K.S. Um, T. Matose, M.M. Tomaszewski, L.K. Barker, M. Mosavel, L.A. Siminoff, H.M. Traino, P. Flicek, T. Juettemann, M. Ruffier, D. Sheppard, K. Taylor, S.J. Trevanion, D.R. Zerbino, B. Craft, M. Goldman, M. Haeussler, W.J. Kent, C.M. Lee, B. Paten, K.R. Rosenbloom, J. Vivian, J. Zhu, D.L. Nicolae, N.J. Cox, H.K. Im, GTEx Consortium, D.A.& C.C. (LDACC)—Analysis W.G. Laboratory, Statistical Methods groups— Analysis Working Group, Enhancing GTEx (eGTEx) groups, NIH Common Fund, NIH/NCI, NIH/NHGrI, NIH/NIMH, NIH/NIDA, Biospecimen Collection Source Site—NDrI, Biospecimen Collection Source Site—rPCI, Biospecimen Core resource—VArI, Brain Bank repository— University of Miami Brain Endowment Bank, Leidos Biomedical—Project Management, ELSI Study, Genome Browser Data Integration & Visualization—EBI, U. of C.S.C. Genome Browser Data Integration & Visualization—UCSC Genomics Institute, Exploring the phenotypic consequences of tissue specific gene expression variation inferred from GWAS summary statistics, Nature Communications 9 (2018) 1825. 10.1038/s41467-018-03621-1.

[42] A. Winter, W. Krämer, F.A.O. Werner, S. Kollers, S. Kata, G. Durstewitz, J. Buitkamp, J.E. Womack, G. Thaller, R. Fries, Association of a lysine-232/alanine polymorphism in a bovine gene encoding acyl-CoA:diacylglycerol acyltransferase (DGAT1) with variation at a quantitative trait locus for milk fat content, Proceedings of the National Academy of Sciences 99 (2002) 9300–9305. 10.1073/pnas.142293799.

[43] W. Ayalew, X. Wu, G.M. Tarekegn, T. Sisay Tessema, R. Naboulsi, R. Van Damme, E. Bongcam-Rudloff, Z. Edea, M. Chu, S. Enquahone, C. Liang, P. Yan, Whole Genome Scan Uncovers Candidate Genes Related to Milk Production Traits in Barka Cattle, International Journal of Molecular Sciences 25 (2024). 10.3390/ijms25116142.

[44] H.G. Olsen, H. Nilsen, B. Hayes, P.R. Berg, M. Svendsen, S. Lien, T. Meuwissen, Genetic support for a quantitative trait nucleotide in the ABCG2 gene affecting milk composition of dairy cattle, BMC Genetics 8 (2007) 32. 10.1186/1471-2156-8-32.

[45] L.-A. Raven, B.G. Cocks, K.E. Kemper, A.J. Chamberlain, C.J. Vander Jagt, M.E. Goddard, B.J. Hayes, Targeted imputation of sequence variants and gene expression profiling identifies twelve candidate genes associated with lactation volume, composition and calving interval in dairy cattle., Mamm Genome 27 (2016) 81–97. 10.1007/s00335-015-9613-8.

[46] H. Pausch, R. Emmerling, B. Gredler-Grandl, R. Fries, H.D. Daetwyler, M.E. Goddard, Meta-analysis of sequence-based association studies across three cattle breeds reveals 25 QTL for fat and protein percentages in milk at nucleotide resolution., BMC Genomics 18 (2017) 853. 10.1186/s12864-017-4263-8.

[47] K.M. Tiplady, T.J. Lopdell, E. Reynolds, R.G. Sherlock, M. Keehan, T.JJ. Johnson, J.E. Pryce, S.R. Davis, R.J. Spelman, B.L. Harris, D.J. Garrick, M.D. Littlejohn, Sequence-based genome-wide association study of individual milk mid-infrared wavenumbers in mixed-breed dairy cattle, Genetics Selection Evolution 53 (2021) 62. 10.1186/s12711-021-00648-9.

[48] H. Atashi, C. Bastin, H. Wilmot, S. Vanderick, X. Hubin, N. Gengler, Genome-wide association study for selected cheese-making properties in Dual-Purpose Belgian Blue cows, Journal of Dairy Science 105 (2022) 8972–8988. 10.3168/jds.2022-21780.

[49] B. Grisart, W. Coppieters, F. Farnir, L. Karim, C. Ford, P. Berzi, N. Cambisano, M. Mni, S. Reid, P. Simon, R. Spelman, M. Georges, R. Snell, Positional Candidate Cloning of a QTL in Dairy Cattle: Identification of a Missense Mutation in the Bovine DGAT1 Gene with Major Effect on Milk Yield and Composition, Genome Research 12 (2002) 222–231. 10.1101/gr.224202.

[50] N.A. Marín-Garzón, A.F.B. Magalhães, P.I. Schmidt, M. Serna, L.F.S. Fonseca, B.M. Salatta, G.B. Frezarim, G.A. Fernandes-Júnior, T. Bresolin, R. Carvalheiro, L.G. Albuquerque, Genome-wide scan reveals genomic regions and candidate genes underlying direct and maternal effects of preweaning calf mortality in Nellore cattle, Genomics 113 (2021) 1386–1395. 10.1016/j.ygeno.2021.02.021.

[51] M. Védrine, F.B. Gilbert, S. Maman, C. Klopp, C. Gitton, P. Rainard, P. Germon, Soluble CD14 produced by bovine mammary epithelial cells modulates their response to full length LPS, Veterinary Research 55 (2024) 76. 10.1186/s13567-024-01329-3.

[52] T.J. Lopdell, K. Tiplady, C. Couldrey, T.J.J. Johnson, M. Keehan, S.R. Davis, B.L. Harris, R.J. Spelman, R.G. Snell, M.D. Littlejohn, Multiple QTL underlie milk phenotypes at the CSF2RB locus., Genet Sel Evol 51 (2019) 3. 10.1186/s12711-019-0446-x.

[53] G. Reales, C. Wallace, Sharing GWAS summary statistics results in more citations, Communications Biology 6 (2023) 116. 10.1038/s42003-023-04497-8.

[54] J.A.L. MacArthur, A. Buniello, L.W. Harris, J. Hayhurst, A. McMahon, E. Sollis, M. Cerezo, P. Hall, E. Lewis, P.L. Whetzel, O.G. Bahcall, I. Barroso, R.J. Carroll, M. Inouye, T.A. Manolio, S.S. Rich, L.A. Hindorff, K. Wiley, H. Parkinson, Workshop proceedings: GWAS summary statistics standards and sharing, Cell Genomics 1 (2021). 10.1016/j.xgen.2021.100004.

[55] A.C. Fernandes, A. Reverter, K. Keogh, P.A. Alexandre, J. Afonso, J.C.P. Palhares, T.F. Cardoso, J.M. Malheiros, J.J. Bruscadin, P.S.N. de Oliveira, G.B. Mourão, L.C. de Almeida Regitano, L.L. Coutinho, Transcriptional response to an alternative diet on liver, muscle, and rumen of beef cattle, Scientific Reports 14 (2024) 13682. 10.1038/s41598-024-63619-2.

[56] P. Di Tommaso, M. Chatzou, E.W. Floden, P.P. Barja, E. Palumbo, C. Notredame, Nextflow enables reproducible computational workflows, Nature Biotechnology 35 (2017) 316–319. 10.1038/nbt.3820.

[57] B.E. Langer, A. Amaral, M.-O. Baudement, F. Bonath, M. Charles, P.K. Chitneedi, E.L. Clark, P. Di Tommaso, S. Djebali, P.A. Ewels, S. Eynard, J.A.F. Yates, D. Fischer, E.W. Floden, S. Foissac, G. Gabernet, M.U. Garcia, G. Gillard, M.K. Gundappa, C. Guyomar, C. Hakkaart, F. Hanssen, P.W. Harrison, M. Hörtenhuber, C. Kurylo, C. Kühn, S. Lagarrigue, D. Lallias, D.J. Macqueen, E. Miller, J. Mir-Pedrol, G.C.M. Moreira, S. Nahnsen, H. Patel, A. Peltzer, F. Pitel, Y. Ramayo-Caldas, M. da Câmara Ribeiro-Dantas, D. Rocha, M. Salavati, A. Sokolov, J. Espinosa-Carrasco, C. Notredame, Empowering bioinformatics communities with Nextflow and nf-core, bioRxiv (2024) 2024.05.10.592912. 10.1101/2024.05.10.592912.

[58] C. Kern, Y. Wang, X. Xu, Z. Pan, M. Halstead, G. Chanthavixay, P. Saelao, S. Waters, R. Xiang, A. Chamberlain, I. Korf, M.E. Delany, H.H. Cheng, J.F. Medrano, A.L. Van Eenennaam, C.K. Tuggle, C. Ernst, P. Flicek, G. Quon, P. Ross, H. Zhou, Functional annotations of three domestic animal genomes provide vital resources for comparative and agricultural research, Nature Communications 12 (2021) 1821. 10.1038/s41467-021-22100-8.

[59] E.L. Clark, A.L. Archibald, H.D. Daetwyler, M.A.M. Groenen, P.W. Harrison, R.D. Houston, C. Kühn, S. Lien, D.J. Macqueen, J.M. Reecy, D. Robledo, M. Watson, C.K. Tuggle, E. Giuffra, From FAANG to fork: application of highly annotated genomes to improve farmed animal production, Genome Biology 21 (2020) 285. 10.1186/s13059-020-02197-8.

[60] E. Giuffra, C.K. Tuggle, Functional Annotation of Animal Genomes (FAANG): Current Achievements and Roadmap, Annual Review of Animal Biosciences 7 (2019) 65–88. 10.1146/annurev-animal-020518-114913.

[61] P.W. Harrison, A. Sokolov, A. Nayak, J. Fan, D. Zerbino, G. Cochrane, P. Flicek, The FAANG Data Portal: Global, Open-Access, “FAIR”, and Richly Validated Genotype to Phenotype Data for High-Quality Functional Annotation of Animal Genomes, Frontiers in Genetics 12 (2021). https://www.frontiersin.org/journals/genetics/articles/10.3389/fgene.2021.639238.

[62] P.W. Harrison, M.R. Amode, O. Austine-Orimoloye, A.G. Azov, M. Barba, I. Barnes, A. Becker, R. Bennett, A. Berry, J. Bhai, S.K. Bhurji, S. Boddu, P.R. Branco Lins, L. Brooks, S.B. Ramaraju, L.I. Campbell, M.C. Martinez, M. Charkhchi, K. Chougule, A. Cockburn, C. Davidson, N.H. De Silva, K. Dodiya, S. Donaldson, B. El Houdaigui, T.E. Naboulsi, R. Fatima, C.G. Giron, T. Genez, D. Grigoriadis, G.S. Ghattaoraya, J.G. Martinez, T.A. Gurbich, M. Hardy, Z. Hollis, T. Hourlier, T. Hunt, M. Kay, V. Kaykala, T. Le, D. Lemos, D. Lodha, D. Marques-Coelho, G. Maslen, G.A. Merino, L.P. Mirabueno, A. Mushtaq, S.N. Hossain, D.N. Ogeh, M.P. Sakthivel, A. Parker, M. Perry, I. Piližota, D. Poppleton, I. Prosovetskaia, S. Raj, J.G. Pérez-Silva, A.I.A. Salam, S. Saraf, N. Saraiva-Agostinho, D. Sheppard, S. Sinha, B. Sipos, V. Sitnik, W. Stark, E. Steed, M.-M. Suner, L. Surapaneni, K. Sutinen, F.F. Tricomi, D. Urbina-Gómez, A. Veidenberg, T.A. Walsh, D. Ware, E. Wass, N.L. Willhoft, J. Allen, J. Alvarez-Jarreta, M. Chakiachvili, B. Flint, S. Giorgetti, L. Haggerty, G.R. Ilsley, J. Keatley, J.E. Loveland, B. Moore, J.M. Mudge, G. Naamati, J. Tate, S.J. Trevanion, A. Winterbottom, A. Frankish, S.E. Hunt, F. Cunningham, S. Dyer, R.D. Finn, F.J. Martin, A.D. Yates, Ensembl 2024., Nucleic Acids Res 52 (2024) D891–D899. 10.1093/nar/gkad1049.

[63] M. Martin, Cutadapt removes adapter sequences from high-throughput sequencing reads, EMBnet.Journal; Vol 17, No 1: Next Generation Sequencing Data AnalysisDO - 10.14806/Ej.17.1.200 (2011). https://journal.embnet.org/index.php/embnetjournal/article/view/200.

