## Supplementary Figures for "A novel reusable transcriptome-wide association study workflow used to map key genes linked to important cattle traits"

### Slide 1
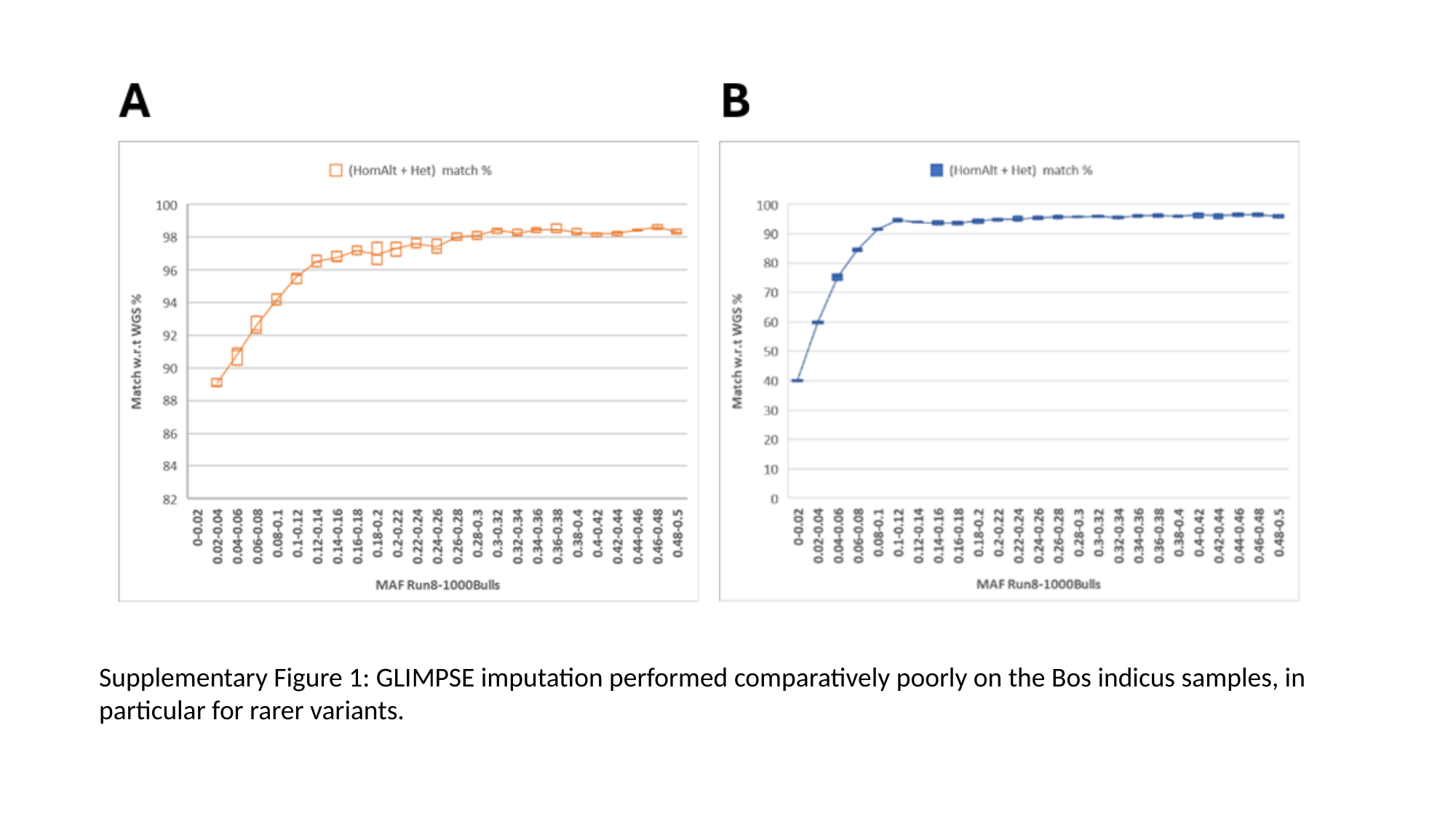

Supplementary Figure 1: GLIMPSE imputation performed comparatively poorly on the Bos indicus samples, in particular for rarer variants.

### Slide 2
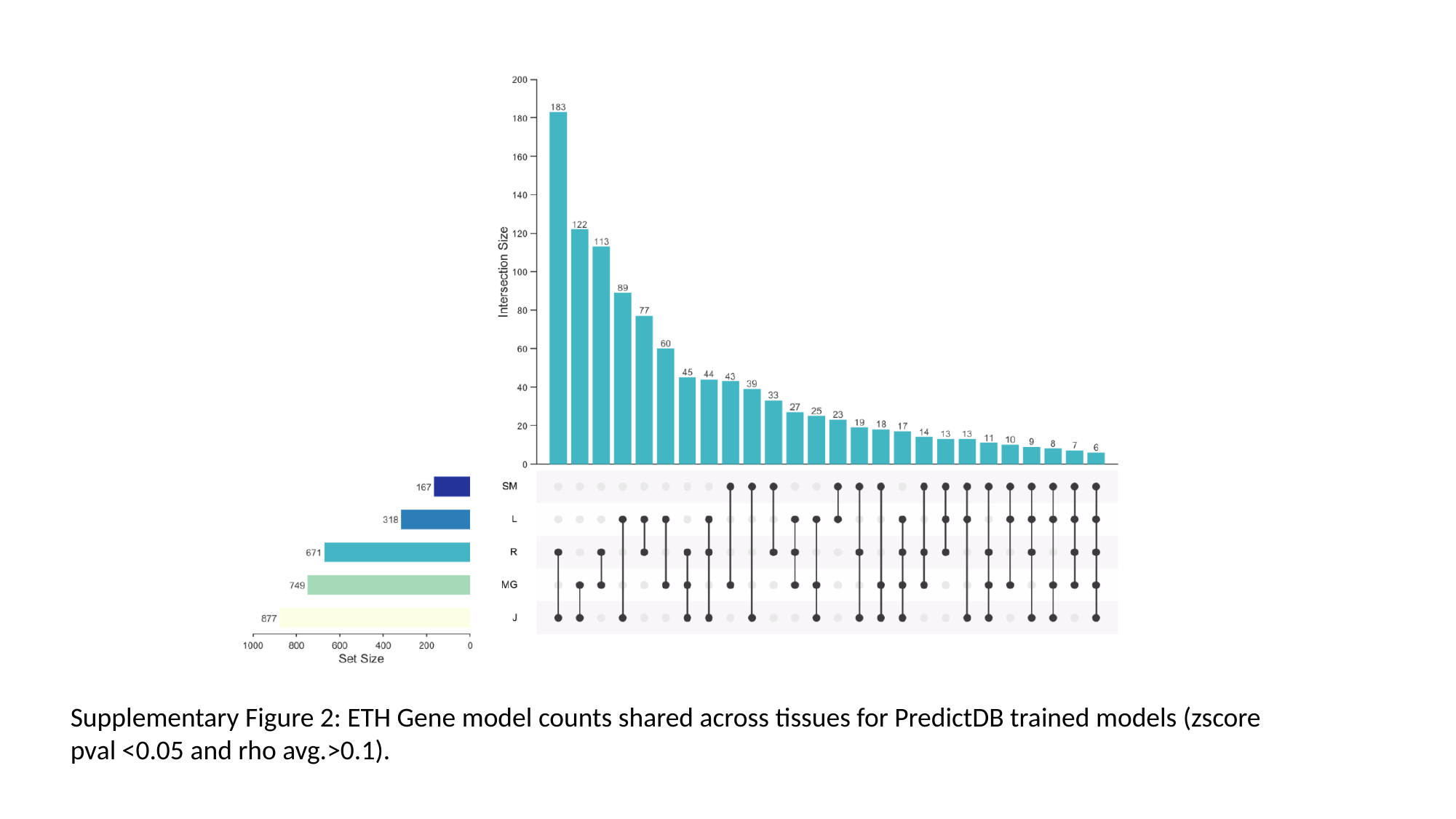

Supplementary Figure 2: ETH Gene model counts shared across tissues for PredictDB trained models (zscore pval <0.05 and rho avg.>0.1).

### Slide 3
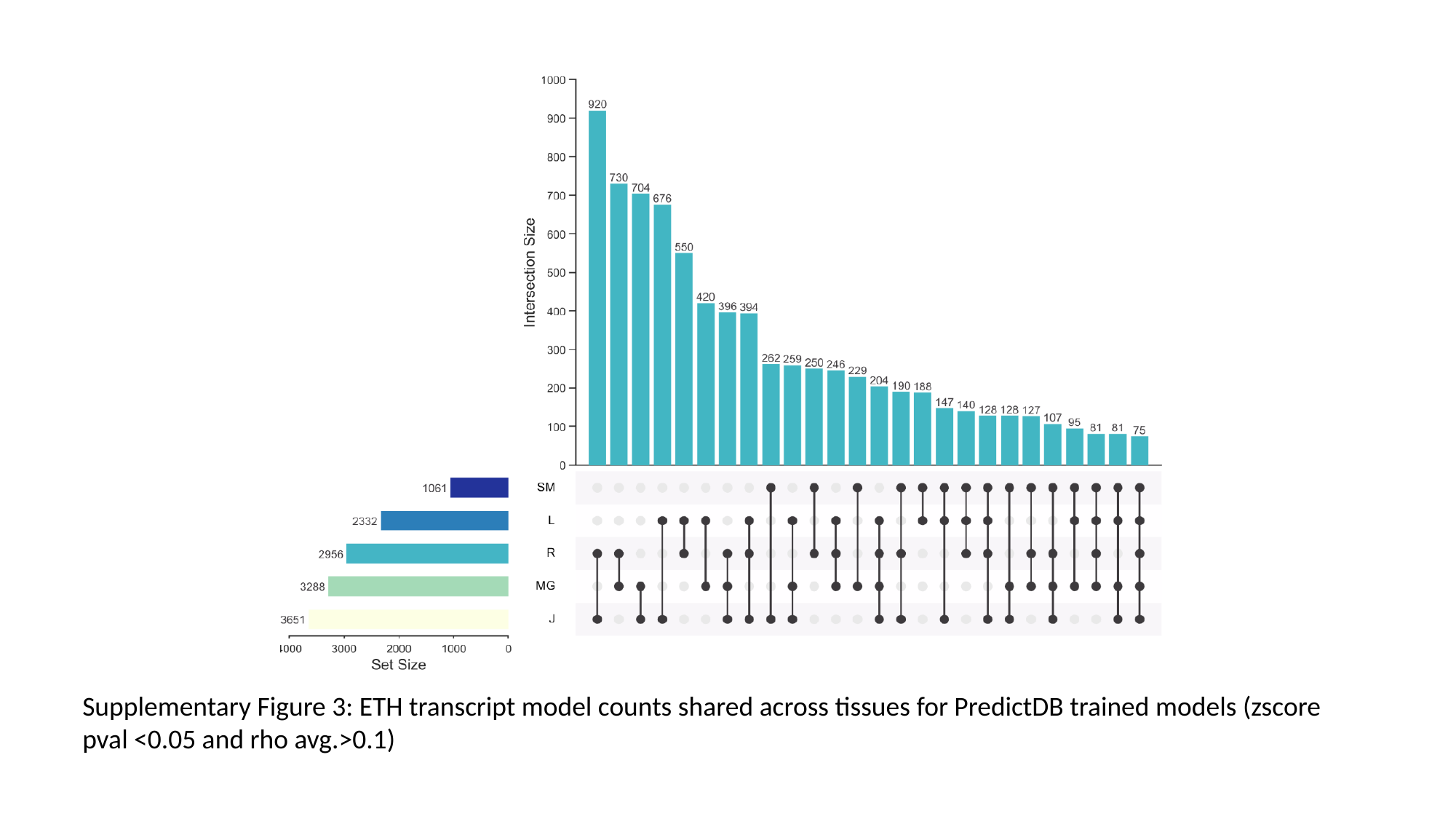

Supplementary Figure 3: ETH transcript model counts shared across tissues for PredictDB trained models (zscore pval <0.05 and rho avg.>0.1)

### Slide 4
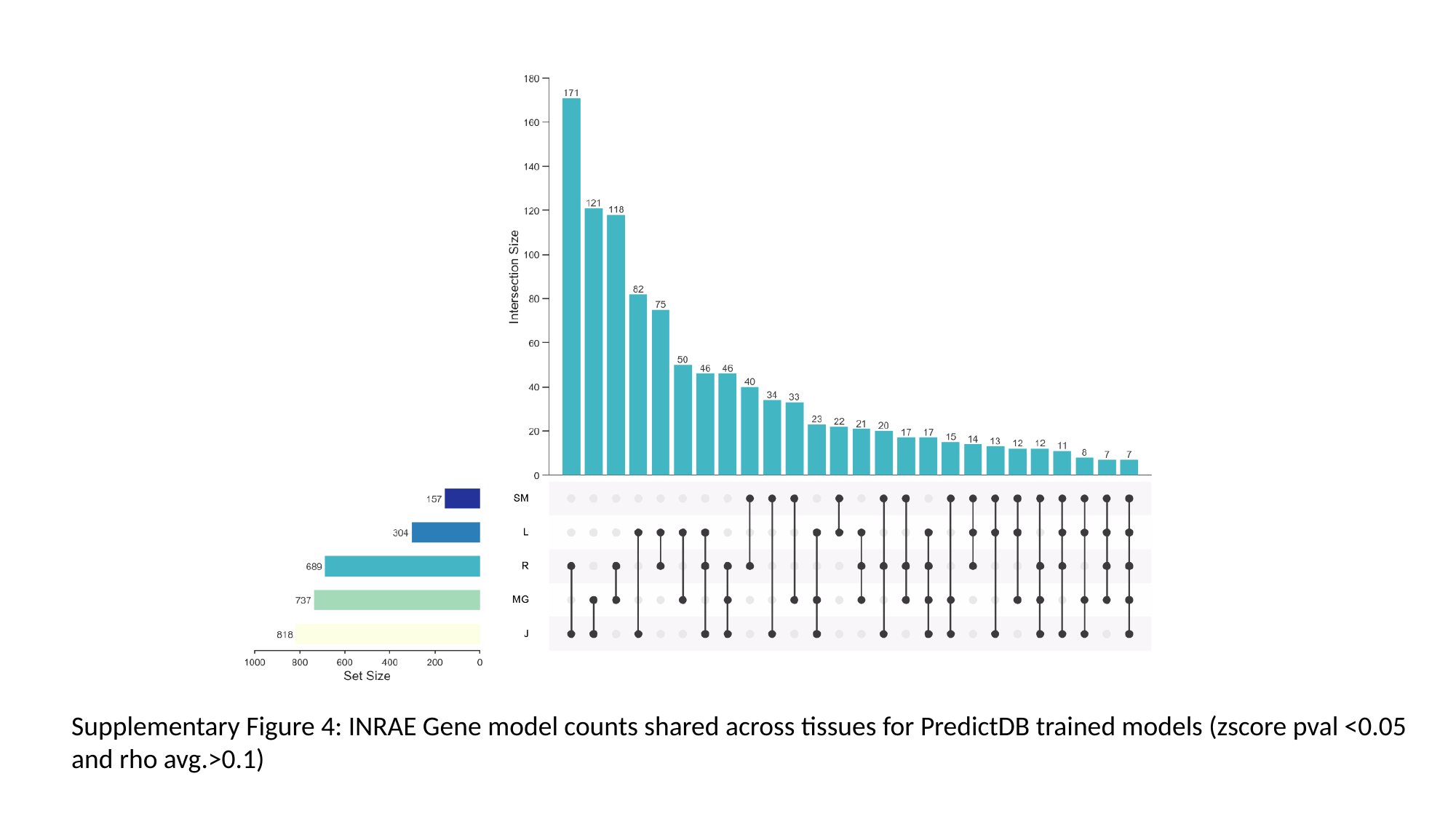

Supplementary Figure 4: INRAE Gene model counts shared across tissues for PredictDB trained models (zscore pval <0.05 and rho avg.>0.1)

### Slide 5
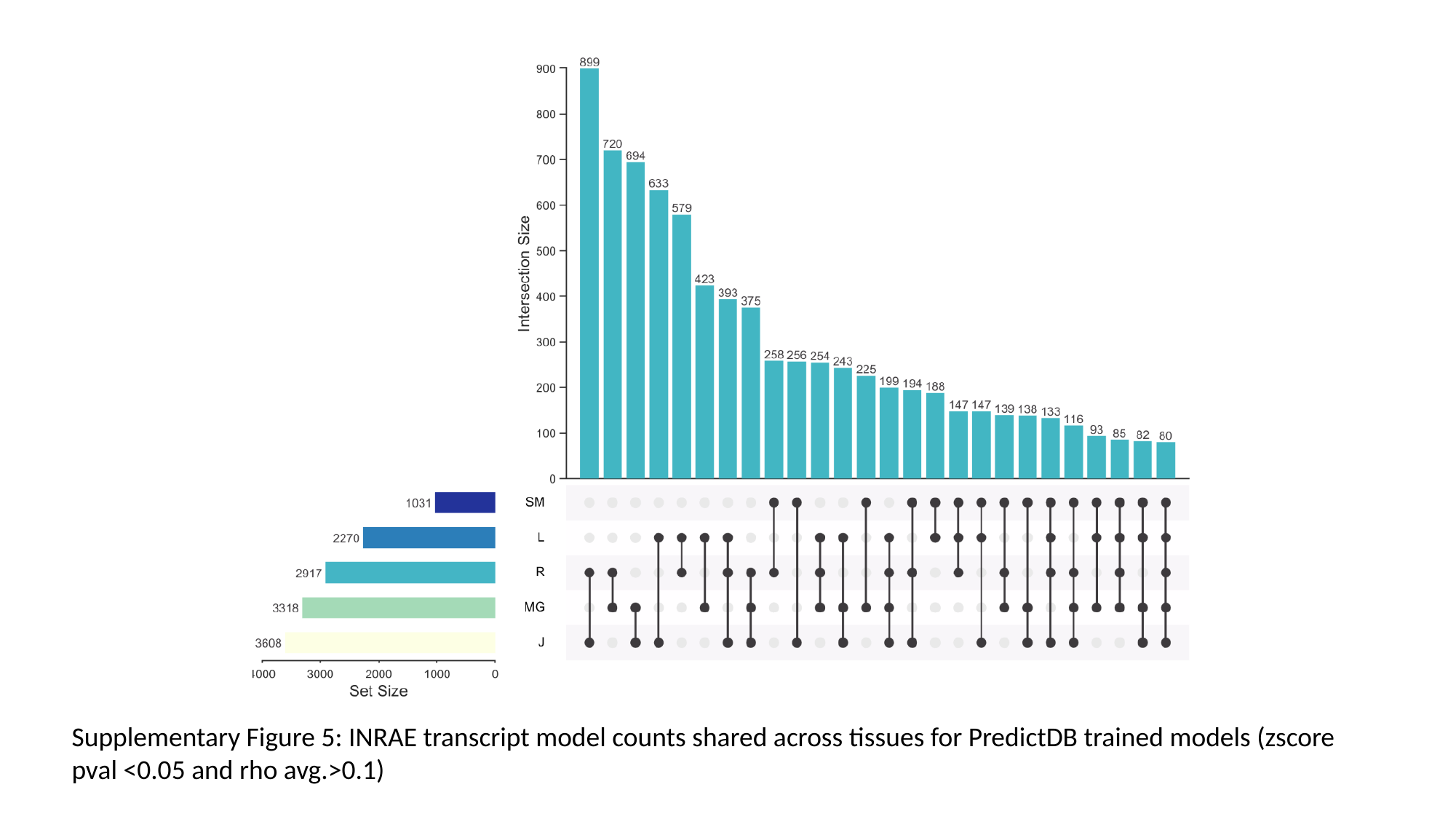

Supplementary Figure 5: INRAE transcript model counts shared across tissues for PredictDB trained models (zscore pval <0.05 and rho avg.>0.1)

### Slide 6
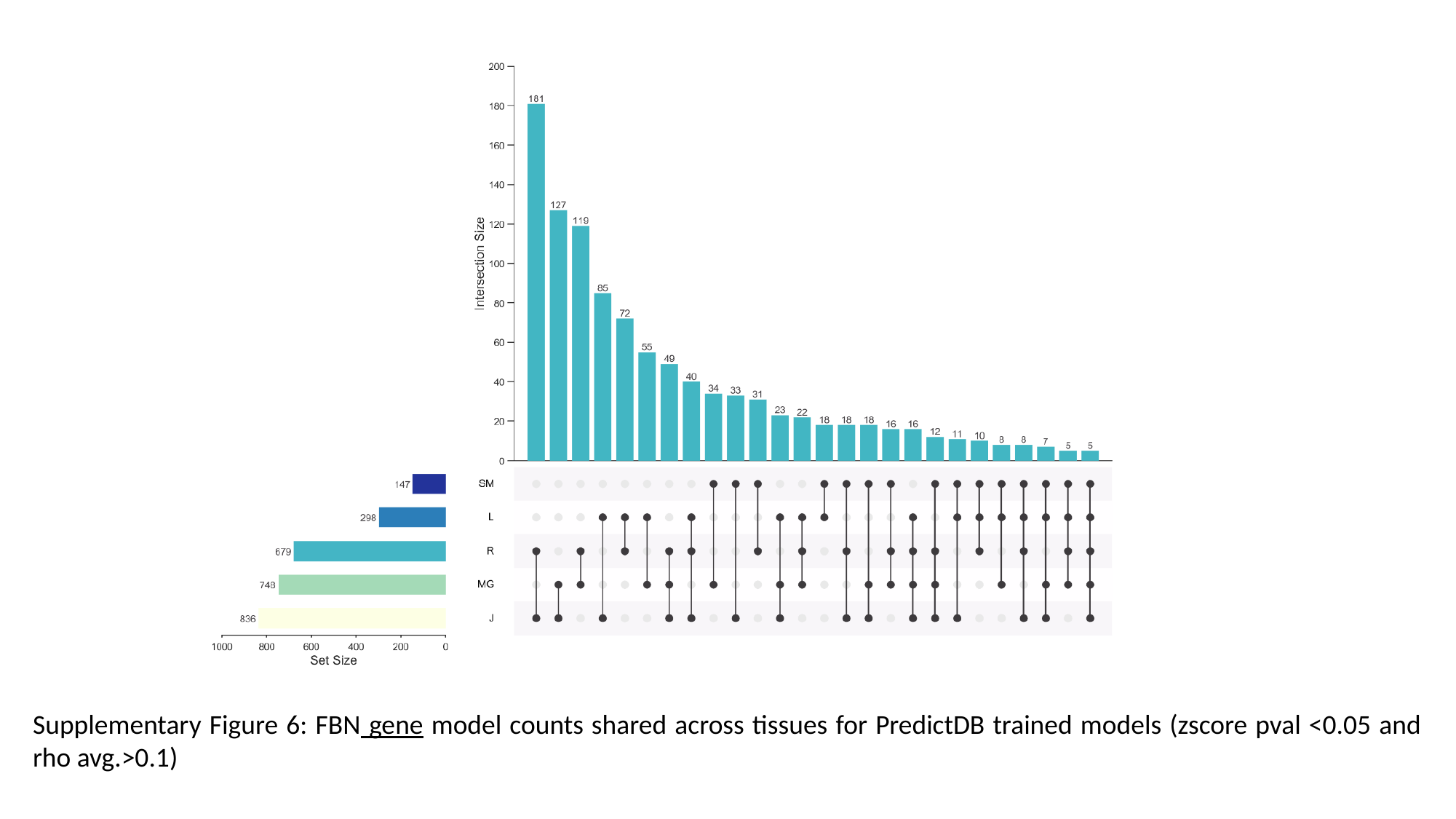

Supplementary Figure 6: FBN gene model counts shared across tissues for PredictDB trained models (zscore pval <0.05 and rho avg.>0.1)

### Slide 7
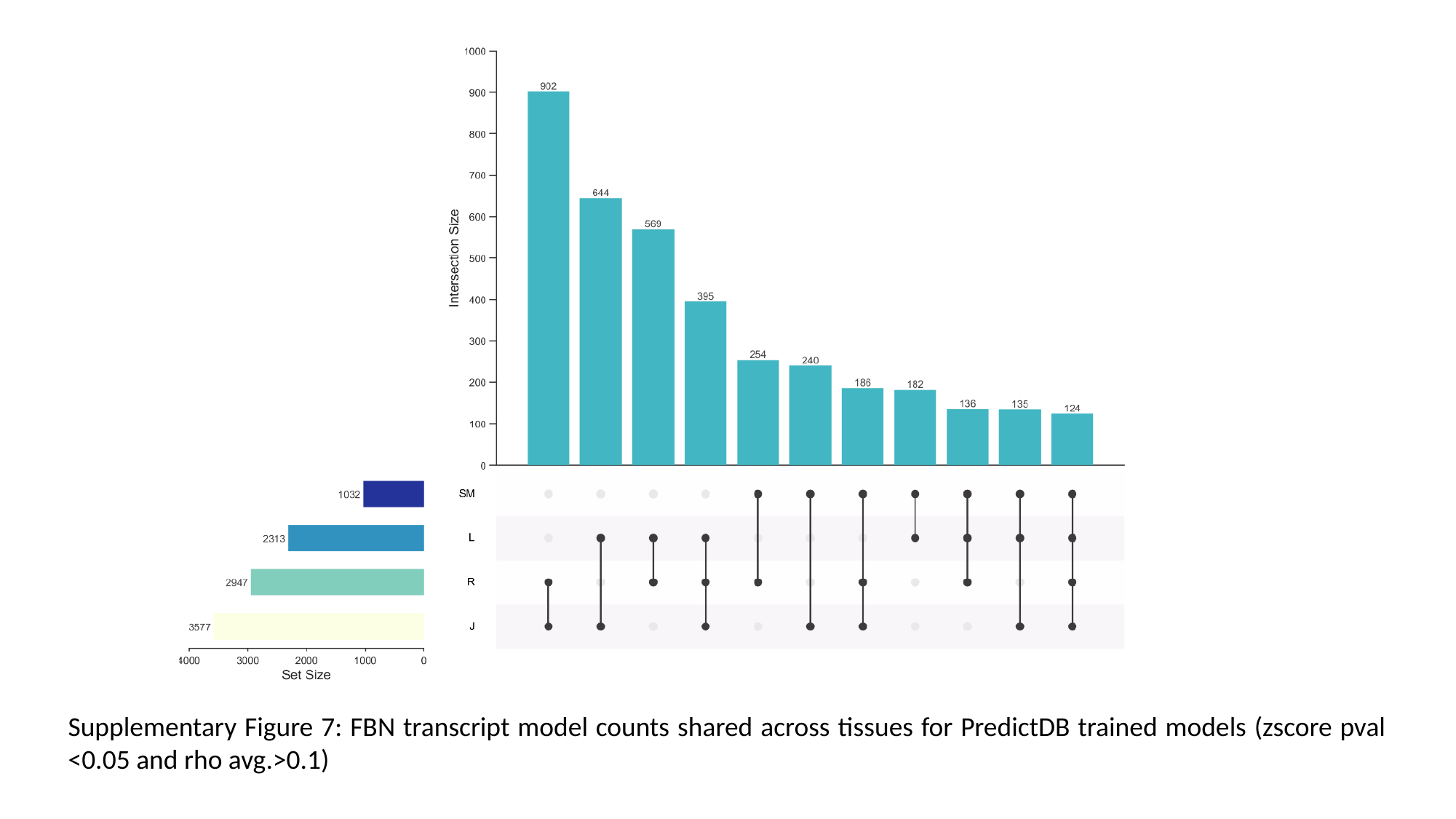

Supplementary Figure 7: FBN transcript model counts shared across tissues for PredictDB trained models (zscore pval <0.05 and rho avg.>0.1)

### Slide 8
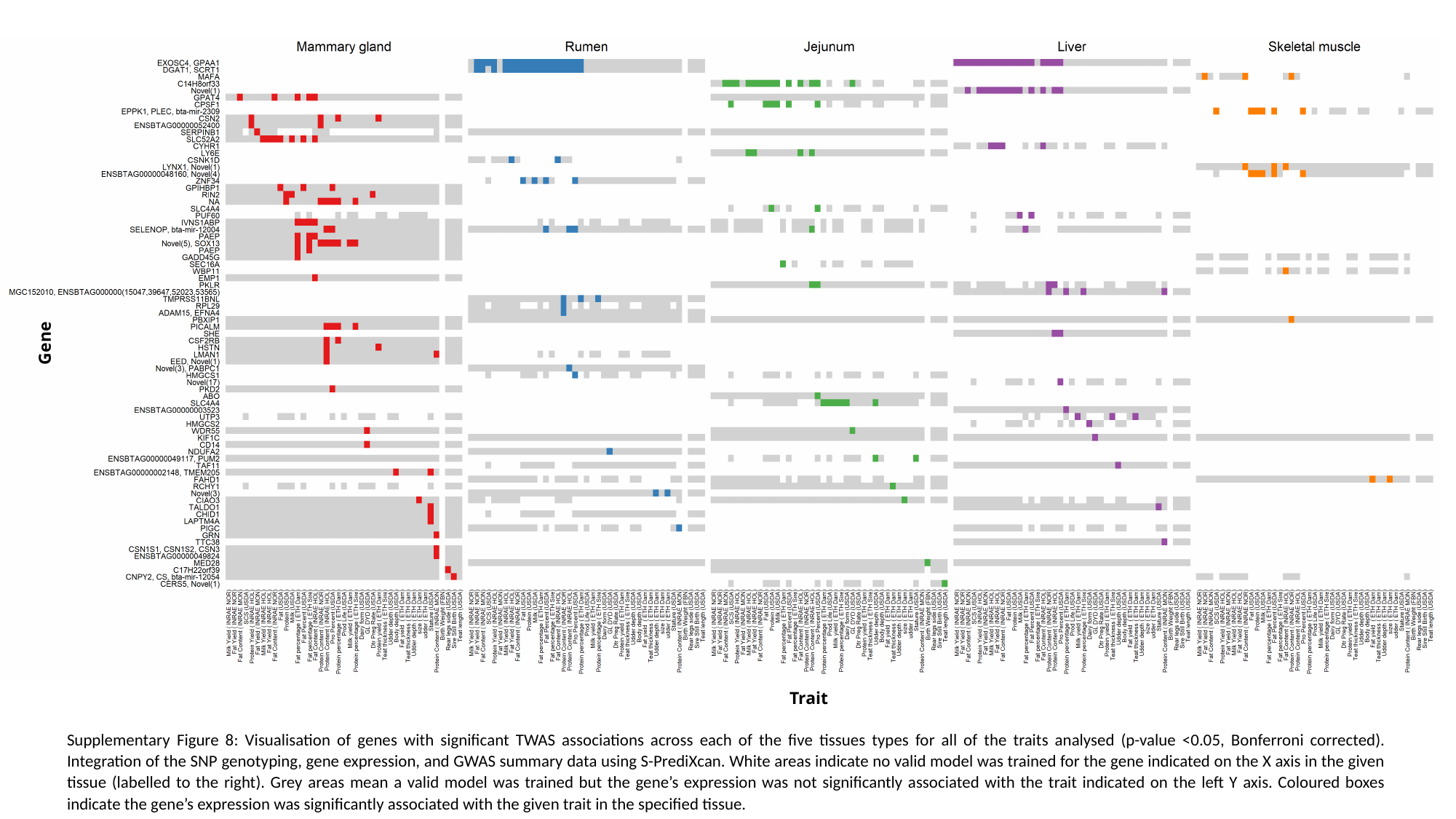

Gene
Trait
Supplementary Figure 8: Visualisation of genes with significant TWAS associations across each of the five tissues types for all of the traits analysed (p-value <0.05, Bonferroni corrected). Integration of the SNP genotyping, gene expression, and GWAS summary data using S-PrediXcan. White areas indicate no valid model was trained for the gene indicated on the X axis in the given tissue (labelled to the right). Grey areas mean a valid model was trained but the gene’s expression was not significantly associated with the trait indicated on the left Y axis. Coloured boxes indicate the gene’s expression was significantly associated with the given trait in the specified tissue.

### Slide 9
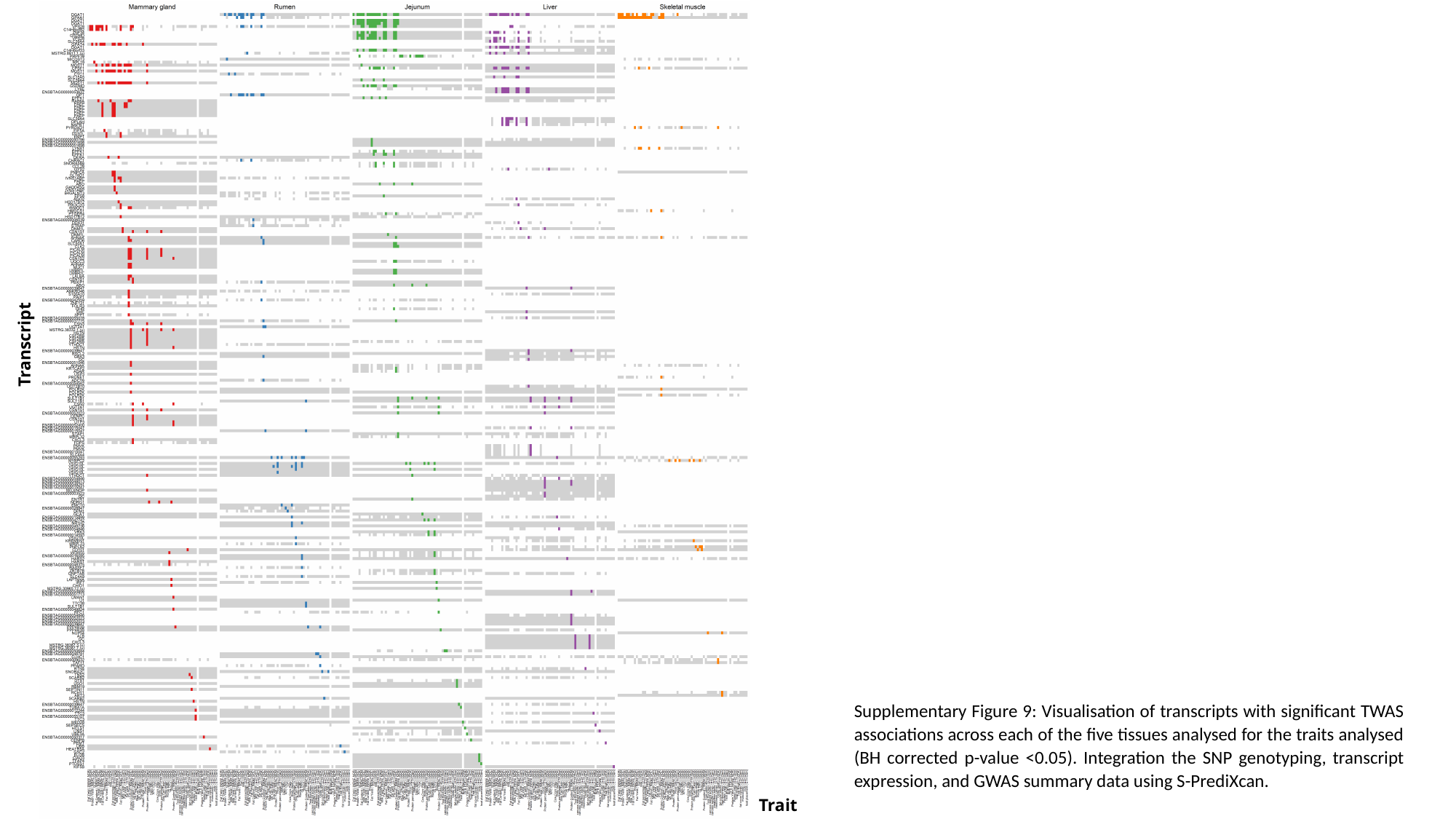

Transcript
Supplementary Figure 9: Visualisation of transcripts with significant TWAS associations across each of the five tissues analysed for the traits analysed (BH corrected p-value <0.05). Integration the SNP genotyping, transcript expression, and GWAS summary data using S-PrediXcan.
Trait

### Slide 10
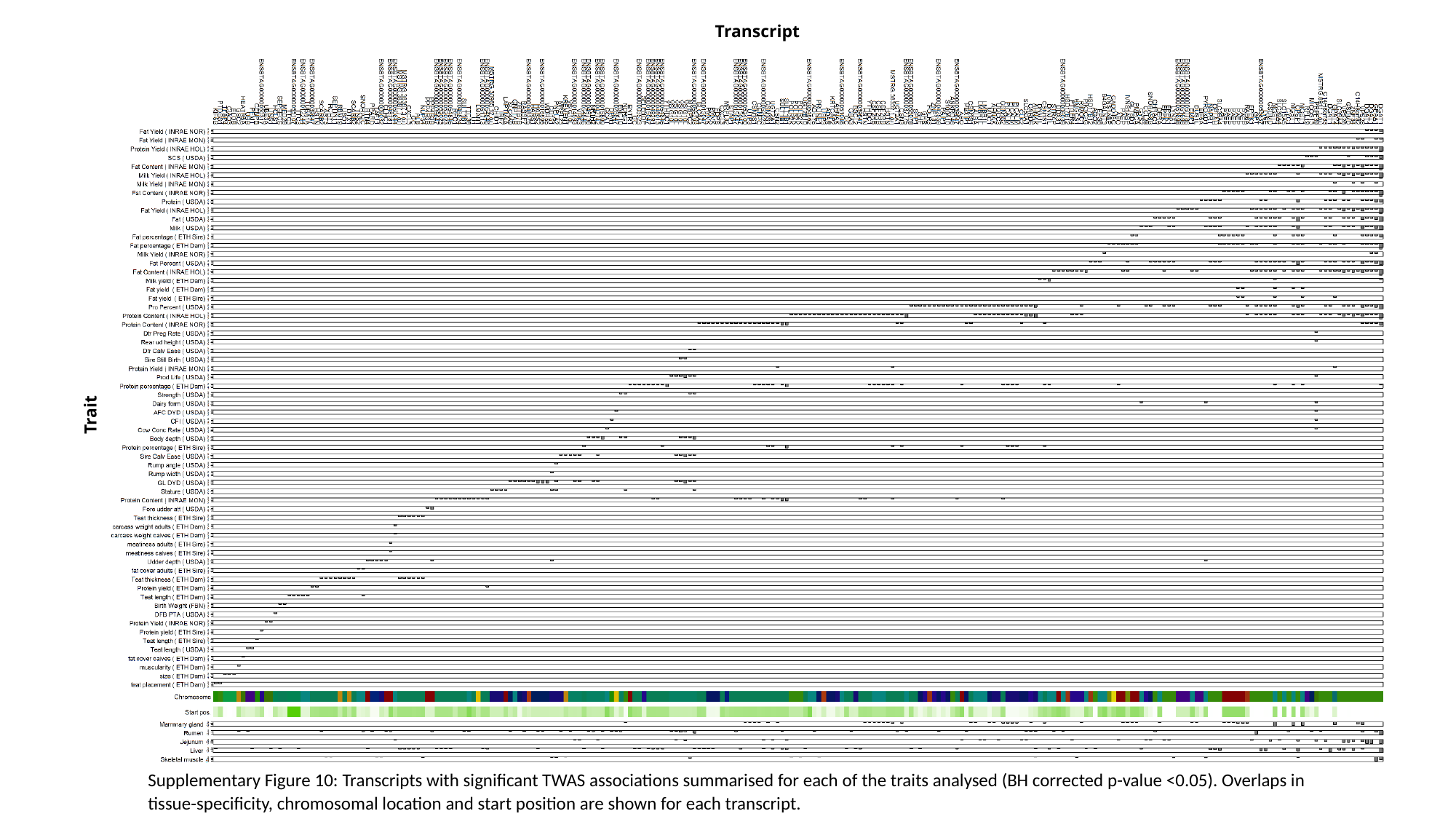

Transcript
Trait
Supplementary Figure 10: Transcripts with significant TWAS associations summarised for each of the traits analysed (BH corrected p-value <0.05). Overlaps in tissue-specificity, chromosomal location and start position are shown for each transcript.
